# State Dependent Beta Oscillations in the Cortico-Basal Ganglia Circuit and their Neuromodulation under Phase-Locked Inputs

**DOI:** 10.1101/2020.03.20.000711

**Authors:** Timothy O. West, Simon F. Farmer, Peter J. Magill, Andrew Sharott, Vladimir Litvak, Hayriye Cagnan

## Abstract

State-of-the-art therapeutic brain stimulation strategies are delivered open loop, using fixed parameters. However, brain states exhibit spontaneous fluctuations dependent upon different behavioural or disease states. Here, we use a model of the cortico-basal ganglia-thalamic circuit to demonstrate how connectivity underpins changes in subcortical beta oscillations – a commonly used control parameter for deep brain stimulation in Parkinson’s disease. We show that recurrent cortical-subcortical loops involving either the cortico-subthalamic or pallido-subthalamic pathways can act in antagonism to modulate the expression of beta band activity (14-30 Hz). These pathways alter the relative timing of intermittent activity across the network, with increased pallido-subthalamic connectivity increasing the propensity of the circuit to enter a state of autonomous oscillation. We demonstrate that phase-locked stimulation can modulate these oscillations, with an efficacy that ultimately depends upon the connectivity across the circuit. This work outlines critical factors required to implement state-adaptive closed-loop brain stimulation.

**Highlights:**

- Converging inputs to the subthalamic nucleus arriving via the external segment of globus pallidus and cortex act in antagonism and promote different beta rhythms.
- Phase locked stimulation has the capacity to selectively enhance or suppress a brain rhythm depending on the stimulation timing.
- The efficacy of stimulation and the parameters required to deliver it, e.g. stimulation timing, effective sensing and stimulation locations, are functions of network state.

## Introduction

The delivery of therapeutic brain stimulation, such as continuous deep brain stimulation (cDBS) for Parkinson’s disease (PD), is typically defined by statically parameterized, open-loop controllers. Stimulation parameters are set in the clinic in order to maximize the amelioration of patient’s symptoms while at the same time minimizing side effects (Volkmann et al., 2009). cDBS suppresses aberrant neural oscillations that emerge during so called ‘oscillopathies’ (Llinás et al., 1999; Brown et al., 2001; Dostrovsky and Bergman, 2004; Uhlhaas and Singer, 2006; Brittain and Brown, 2014). For example, in dopamine depleted states associated with PD, oscillatory activity in the beta frequencies (14-30 Hz), normally engaged during the functional control of movement (Engel and Fries, 2010; Jenkinson and Brown, 2011; Khanna and Carmena, 2015; Palmer et al., 2016), becomes abnormally amplified across the circuits formed by the cortex, thalamus, and basal ganglia (Brown et al. 2001; Levy 2002; Mallet et al. 2008b; Weinberger et al. 2006). These pathological rhythms are widely synchronized with their frequency and amplitude modulated by specific brain states such as rest, movement, anaesthesia, and changes in dopamine levels (Sharott et al., 2005a; de Solages et al., 2010; Litvak et al., 2011; van Wijk et al., 2012; Brazhnik et al., 2014; Delaville et al., 2014; West et al., 2016, 2018).

Neural activity in the beta frequencies exhibits intermittent fluctuations, consisting of high-amplitude sub-second events. These are referred to as ‘bursts’ and have been characterised in both pathological (Tinkhauser et al., 2017b; Torrecillos et al., 2018; Cagnan et al., 2019b) and functional (Feingold et al., 2015; Lundqvist et al., 2016; Sherman et al., 2016; Khanna and Carmena, 2017; Shin et al., 2017; Little et al., 2019) brain activity. Recent experiments have demonstrated that these rhythms can be specifically interrupted using adaptive DBS (aDBS; Little et al., 2013; Arlotti et al., 2016; Tinkhauser et al., 2017a) – a closed-loop control strategy that triggers stimulation at the onset of a beta burst. Importantly the delivery of stimulation in the context of ongoing neural activity has important impacts upon behaviour (Stagg et al., 2011; Pirulli et al., 2013; Martin et al., 2014; Herz et al., 2018; Kahan et al., 2019) indicative of state dependent efficacy (Silvanto et al., 2008; Kahan et al., 2019). It is thus important to understand how the precise timing of stimulating input interacts with ongoing, spontaneous fluctuations in neural activity (McIntyre and Hahn, 2010; Cagnan et al., 2019a).

To understand how the delivery of stimulation can be optimized to interact with state dependent fluctuations in rhythms such as Parkinsonian beta bursts, it is necessary to derive the relationship between the structural connectivity of the circuits responsible for their generation, propagation, and maintenance. Several candidate circuits have been proposed, these include: the pallido-subthalamic feedback loop (Bevan et al., 2002; Terman et al., 2002; Cruz et al., 2011; Tachibana et al., 2011; Liu et al., 2017; Shouno et al., 2017); cortico-basal ganglia thalamic ‘long loop’ (Pavlides et al., 2015; Brazhnik et al., 2016); the thalamocortical relay (van Albada et al., 2009; Reis et al., 2019); striato-pallidal feedback (Corbit et al., 2016; Blenkinsop et al., 2017; Crompe et al., 2020); changes in striatal outflow (Gillies and Willshaw, 2004; Brown, 2007; Hammond et al., 2007; Kumar et al., 2011; Sharott et al., 2017); and intrinsic striatal dynamics (McCarthy et al., 2011). It is likely that the observed activity arises from a competition of the output of these circuits (Leblois et al., 2006; Pavlides et al., 2015; Fountas and Shanahan, 2017).

Propagation of beta activity is accompanied by altered phase synchronization of neural activity across the cortico-basal ganglia circuit (Cagnan et al., 2015, 2019b). Novel therapies aim to deliver stimulation that is phase-locked to an ongoing control signal such as the peripheral tremor amplitude (Brittain et al., 2013; Cagnan et al., 2017) or neuronal beta activity (Holt et al., 2019; Peles et al., 2020; Sanabria et al., 2020) in order to specifically target phase synchronization across circuits. Phase-locked stimulation has also been shown to modulate coherent neural activity underpinning healthy communication in the brain and modulate behaviour (Polanía et al., 2012; Siegle and Wilson, 2014; Cordon et al., 2018; Zanos et al., 2018).

In this work we use a computational model of the cortico-basal ganglia-thalamic circuit (Moran et al., 2011; Marreiros et al., 2013; van Wijk et al., 2018) that has been fit (West et al., 2019) to *in vivo* intracranial recordings of neuronal activity obtained from rats rendered Parkinsonian by chronic dopamine depletion (Mallet et al., 2008b). This model demonstrates that rhythmic activity within this circuit is the manifestation of an interplay between reciprocal cortico-basal ganglia loops. Furthermore, these rhythms can be manipulated using precisely timed, phase-locked stimulation of the cerebral cortex or STN. We investigate the interaction between state-dependent changes in the expression of beta rhythms and the efficacy of phase-locked stimulation to better inform how adaptive control of stimulation can provide optimal therapeutic effects in the face of spontaneous alterations of brain networks.

## Materials and Methods

### Model Optimization with Approximate Bayesian Computation

This study reports a computational model of the population activity within the cortical-basal ganglia-thalamic circuit. In a previous paper, we described a model optimization and model selection procedure based upon the Approximate Bayesian Computation framework (ABC) (West et al. 2019a). This procedure was used to fit 13 different models of the network to empirical data (Magill et al., 2006; Mallet et al., 2008b, 2008a; Moran et al., 2011)., and then subsequently select the candidate model that could best explain its spectral features (see below). For a full formulation of the fitting and model selection procedure please see West et al. (2019a). ABC provides a likelihood free optimization algorithm (Beaumont et al., 2002; Liepe et al., 2014) that is founded in the generation of ‘pseudo-data’ from numerical simulations of a model that can then be compared to the empirical data using a common set of features (e.g. spectral power and directed functional connectivity). By iteratively minimizing the error between a set of data features common to the empirical data and pseudodata through the optimisation of model parameters, the algorithm may converge to an approximation to the distribution over parameters of the model that best explain the observed data. ABC allows for models of steady state features such as spectra to be modelled without any requirements to approximate the system’s behaviour by its equilibria. This leaves us free to explore non-steady state activity, such as transient burst like behaviour. Here, we placed a restriction on the model space to only those yielding intermittent, burst like behaviour. Therefore, models that yielded either exponentially divergent, or highly autocorrelated envelopes were excluded (West et al., 2019) enabling us to explore transient dynamics such as beta bursts in post-hoc numerical simulations of the fitted models. In this study, the population’s local oscillatory profile (i.e. the power spectra) plus the patterns of connectivity between them (i.e. the directed functional connectivity) were used as data features to fit model parameters. To determine directed functional connectivity, we used non-parametric directionality (NPD; see section below).

### Model Description

We employ a model formalism that is based upon a description of the propagation of activity across the neural populations of the cortico-basal ganglia-thalamic network. Specifically, we use a system of coupled neural mass equations (Jansen and Rit, 1995; David and Friston, 2003) to simulate the distributed activity across this network. The biological and theoretical basis of this model has been described previously (Moran et al., 2011; Marreiros et al., 2013; van Wijk et al., 2018). Furthermore, we adapt the original state equations to incorporate stochastic inputs and explicit, finite transmission delays. For the full state space description of the model and technical details regarding the numerical schemes used for integration of the equations, please see West et al. (2019). Overall, the model consists of nine coupled 2^nd^ order stochastic differential equations, each modelling a separate (assumingly homogenous) population of neurons within the circuit. This includes a model of the motor cortex microcircuit consisting of three pyramidal layers (superficial, middle, and deep) plus an inhibitory interneuron population (Bhatt et al., 2016). Each cortical layer also contained recurrent self-inhibitory connections reflecting local neuronal gain control. Furthermore, we modelled neuronal populations in the striatum (STR); the external and internal segments of the globus pallidus (GPe/i); the subthalamic nucleus (STN); and the thalamus (Thal.). The GPi and Thalamus were included as ‘hidden sources’ to model re-entrant feedback from basal ganglia to the cortex. Stochastic inputs (Gaussian noise process with a parameterized gain factor) were delivered to all subcortical populations and the middle pyramidal layer of the motor cortex.

In this paper we bring forward the posterior model fit (specifically – the *maximum a posteriori* estimates of the parameters) from West et al. (2019a) where we used ABC to fit 13 variations of the full model that included or excluded different key connections such as the subthalamo-pallidal connection (STN → GPe) and the hyperdirect pathway (M2 → STN). We then used a Bayesian model comparison approach to determine a subset of these models as the best explanations of the empirical data, accounting for both the individual model’s ability to accurately reproduce the observed data and the extent to which model parameters deviate from a prior distribution.

After discounting for difference in the parsimony of posterior model fits (i.e. the divergence between posterior and prior distributions over parameters), we found that a model incorporating both the hyperdirect and subthalamo-pallidal pathways was the best candidate in describing the patterns of neuronal activity in recordings made in Parkinsonian rats (see below). This posterior model fit is used for the simulations in this paper which we refer to as the *fitted* model and its structure is presented in figure 1.

**Figure 1.**
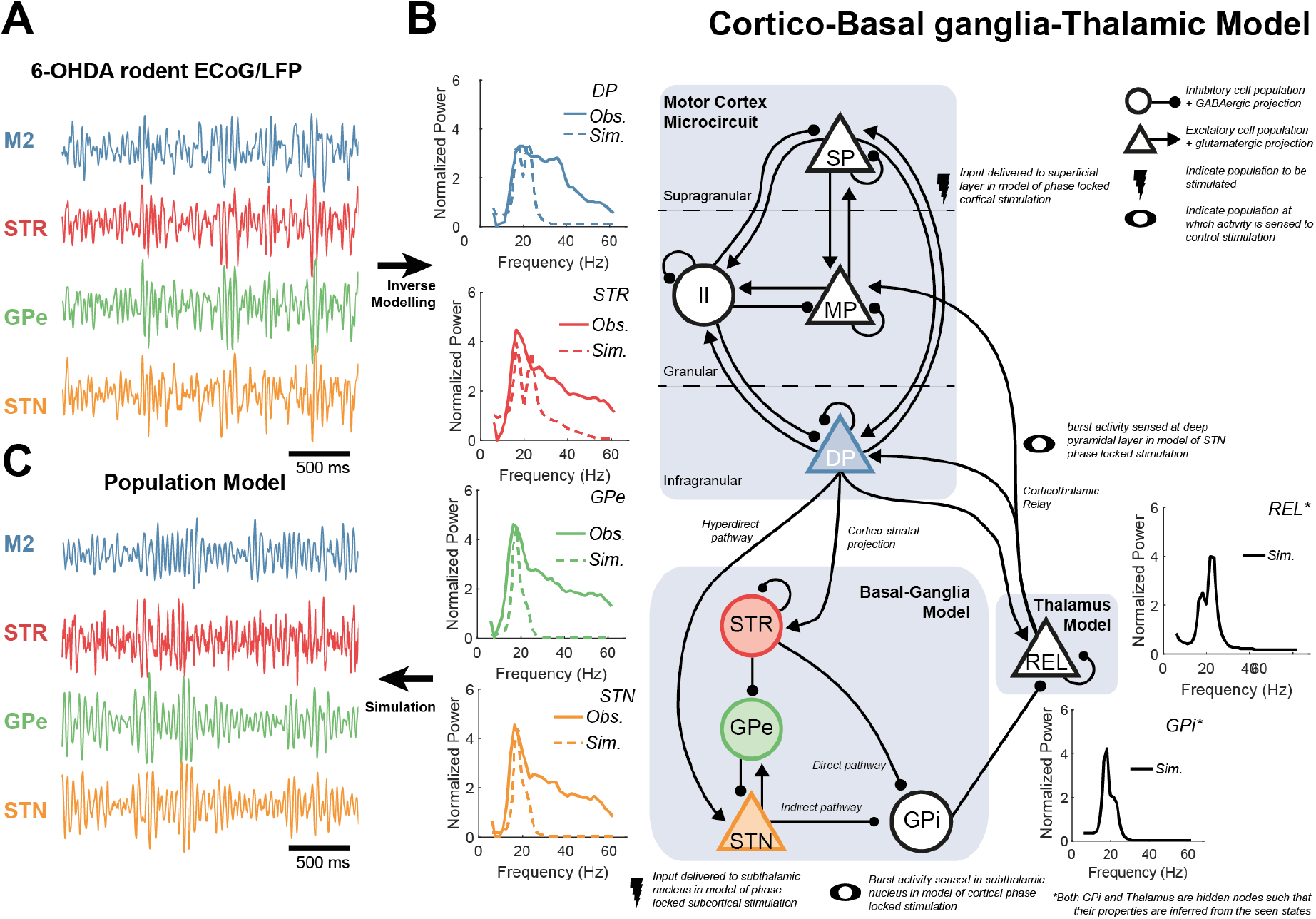
Schematic of cortico-basal ganglia-thalamic model and fit to empirical data from Parkinsonian rodents. A model describing the population activity in this circuit was fit to data features (power spectra and directed functional connectivity) of **(A)** electrophysiological recordings: electrocorticography from motor cortex M2 (blue) as well as local field potentials from striatum STR (red); external segment of the globus pallidus GPe (green); and the subthalamic nucleus STN (yellow) made in a 6-OHDA-lesioned rat model of Parkinsonism (procedure detailed in West et al. 2019a). Data was normalized and band-passed 4-100 Hz before being transformed to the data features used to fit parameters of the computational model. **(B)** Schematic of model architecture, detailing excitatory populations and their glutamatergic projections (triangular nodes with arrows) and inhibitory/GABAergic projections (circular nodes with ball ended arrows). The motor cortex microcircuit (Bhatt et al., 2016) comprises three layers: superficial pyramidal cells (SP; supragranular); middle pyramidal (MP; granular); and deep pyramidal cells (DP; infragranular), plus an inhibitory interneuron population (II). The basal ganglia model comprises four populations, with each node representing activity in the STR, GPe, STN, and internal segment of the pallidus (GPi). The GPi forms the output of the basal ganglia and acts to inhibit relay cells of the thalamus (REL). The main subcortical pathways include the direct, indirect, hyperdirect, and cortico-thalamic interactions. The inset graphs indicate the empirical and simulated power spectra in bold and dashed lines, respectively. For the full set of empirical and fitted data features please see supplementary information I. GPi and REL were treated as hidden nodes and their respective neural activities were inferred from the dynamics of the empirically recorded brain regions. **(C)** Simulations of this circuit yields time series with transient, burst like behaviour similar to that seen *in vivo* (A).

### Empirical Data – 6-hydroxydopamine (6-OHDA)-Lesioned Rats

The models were fitted to archival data consisting of multisite recordings in the basal ganglia and cerebral cortex of nine adult male Sprague-Dawley rats (Charles River, Margate, UK) with 6-OHDA-induced dopamine depletion, a model of the dopaminergic degeneration associated with Parkinsonism in humans as described previously (Magill et al., 2004, 2006). All experiments were conducted in accordance with the Animals (Scientific Procedures) Act, 1986 (United Kingdom), and with Society for Neuroscience Policies on the Use of Animals in Neuroscience Research. Animals were implanted with two multi-contact silicon probes to measure local field potentials (LFP) from multiple structures in the basal ganglia: dorsal striatum, GPe, and STN. Additionally, electrocorticography (ECoG) was measured over “secondary motor cortex” (M2) a homologue of the premotor cortex in humans (Paxinos and Watson, 2007) using a 1 mm diameter steel screw juxtaposed to the dura mater above the right frontal cortex. Anaesthesia was induced with 4% v/v isoflurane (Isoflo, Schering-Plough Ltd., Welwyn Garden City, UK) in O2 and maintained with urethane (1.3 g/kg, i.p.; ethyl carbamate, Sigma, Poole, UK), and supplemental doses of ketamine (30 mg/kg; Ketaset, Willows Francis, Crawley, UK) and xylazine (3 mg/kg; Rompun, Bayer, Germany). Recordings were made during periods of ‘cortical activation’ (Steriade, 2000) induced by a hind-paw pinch. For more details of the experimental recordings and their acquisition please see the original experimental papers (Magill et al., 2006; Mallet et al., 2008b, 2008a; Moran et al., 2011).

Preprocessing of time series data (both LFP and ECoG) were done as follows: all data 1) were down sampled from the hardware native 17.9 kHz to 250 Hz using Spike2 acquisition and analysis software (Cambridge Electronic Design Ltd., Cambridge, UK); 2) imported into MATLAB; 3) mean subtracted; 4) band-passed filtered 4-100 Hz with a finite impulse response, two-pass (zero-lag) filter with order optimized for data length; 5) Z-scored to standardize to unit variance; 6) epoched to 1 second segments; and 7) subjected to a Z-score threshold criterion such that epochs containing any high amplitude artefacts were removed. The exact threshold was chosen on a case by case basis dependent upon recording gain and size of the artefact. This artefact-rejected, epoched data was then taken forward to compute the autospectra and NPD used as features for the ABC fitting procedure.

### Spectral Estimation of Empirical and Simulated Data

Analysis of simulated and empirical data involved computation of power spectra. Power spectra were constructed using the averaged periodogram method across 1 second epochs multiplied by a Hanning window of the same length. Spectra from the nine rats were summarised using the group average. To account for this, the 1/f background was removed by first performing a linear regression on the log-log empirical spectra and then subtracting the linear component from the spectra. This ensured that the parameter estimation scheme was focused upon fitting the spectral peaks in the empirical data and not the profile of 1/f background noise.

To compute functional connectivity (magnitude squared coherence; used only in the analysis of the post-hoc simulations) and directed functional connectivity (NPD; used only as a data feature in the ABC fitting), we used the Neurospec toolbox (http://www.neurospec.org/). NPD provides a non-parametric assessment of directed connections between neural signals derived from their spectral estimates alone (Halliday et al., 2016; West et al., 2020). The resulting NPD spectra were averaged across the nine rats. Features in the group averaged empirical data were smoothed using a sum of Gaussians (maximum of three), with order optimized using adjusted R^2^ to aid the fitting procedures.

### In-Silico Inactivation Experiments to Determine Individual Contribution of Connections

In the first set of simulations we perform a systematic inactivation analysis to determine which connections are required for the generation and/or propagation of beta oscillations in the STN. This was done by taking the fitted model and removing individual connections from the model by setting their weights to zero. We then re-simulate the data and compare the resulting power/magnitude and frequency of the power and coherence with that of the original model in order to determine the percentage change in beta amplitude. We define the power as the sum within the beta band from 14-30 Hz, divided by the number of frequency bins; magnitude as the peak value within this band; and frequency as the location of the peak within this band.

### Impact of Connection Strength upon Rhythmic Activity

In order to determine the influence of not only inactivation but also modulation of connections we also performed an analysis of spectral changes associated with a range of connection strengths. We simulate models with strengths of connections initially ranging from 10% to 1000% of the parameter inferred from the empirical data. The connection strength of the *n^th^* connection *C_n_* is given by:

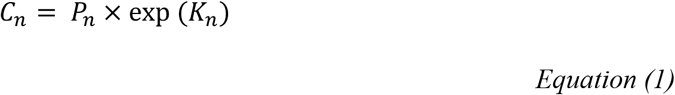

with *P_n_* as the expected value of the posteriors of the fitted model, and *K_n_* as a log scaling parameter. Individual connection strengths are reported as a percentage of the parameter *C_n_* in the fitted model. In the remaining analyses we choose a range of connection strengths from the smallest connection strength examined (10%) up to a maximum strength that yields a 200% increase in the STN beta power.

### Analysis of Recurrent Circuit Activity in the Maintenance of Beta Power and Cortico-Subthalamic Beta Coherence

To investigate how differences in the strengths of loops within the circuit impact the steady state (i.e. the time averaged) properties of beta synchrony within the network, we simulated models with randomly configured connectivity. To this end, we drew 500 sets of connectivity parameters from the posterior distribution (see supplementary table I for values). In order to increase the range of connectivity that was analysed the spread of the posterior distributions was artificially inflated prior to this draw by scaling their variances to be 5 times larger. All other parameters were held fixed at the values specified by the expected value of the posterior derived from the model fit. For each random model, 30 seconds of data was simulated, and the spectral properties were estimated (see Spectral Estimation of Empirical and Simulated Data). To summarise the connectivity within each model we derive two measures:

1. A measure of the sum of absolute connection strength (disregarding the sign of the connection):

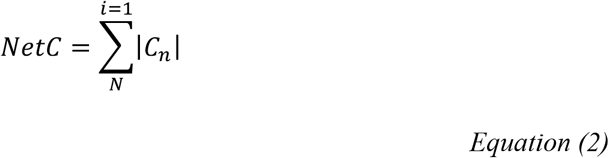

where *C_n_* is the connection strength of the *n^th^* connection. This is then reported as a percentage difference from the mean of the simulated set. This measures the total degree of connectivity within the loop irrespective of whether it comprises inhibitory or excitatory connections. We refer to this as *net loop strength*.
2. In order to measure differences in circuitry arising from either a net increase in excitation or inhibition, we also derive a measure of the E/I balance within a loop:

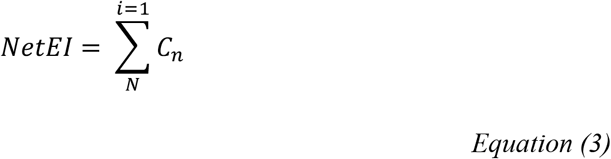 We report this as a difference from the mean of the simulated set. By accounting for the sign of the connection this measure yields the net change in the relative excitation or inhibition within the loop. We refer to this as *net loop E/I balance.*

Using these two measures we then estimate how both change between models yielding either the 1^st^ and 4^th^ quartiles of STN, M2, and STN/M2 coherence at beta (14-30 Hz). Differences in loop strength associated with these two model sets are estimated using Student’s t-test, and statistical significance established using a Bonferroni corrected significance threshold. We also test for the correlation of loop connectivity measures (1) and (2) with power and coherence. In the cases that the non-parametric (Spearman’s) correlation test reached (Bonferroni corrected) significance level, we computed linear regressions of the relations.

### Definition of Burst Events and their Timings

To define and characterize the properties of intermittencies in rhythms within the simulated data, we follow a similar approach to that taken in previous experimental work (Tinkhauser et al., 2017b; Cagnan et al., 2019b) that constructs a band filtered signal and uses a threshold to define burst events as suprathreshold amplitude activity. In this case we use a two-pass, Butterworth filter with a passband of 18± 7.5 Hz. We take the Hilbert transform of this narrowband signal to compute the analytic signal. The amplitude envelope of this signal is then taken as the absolute value of the transform. From this signal we can then define a threshold to determine burst events. In this application we use the 75^th^ percentile of the data and only consider bursts which had a duration longer than at least 1 cycle of an 18 Hz oscillation (middle of the passband) equivalent to 56 ms. In analyses comparing bursts across connection strengths or during phase-locked stimulation, we compute a common threshold that is derived from the 75^th^ percentile of the envelope obtained from the unstimulated, fitted model (i.e. at 100% connectivity). Burst properties such as the median interburst interval (time from offset to onset); the burst amplitude (75^th^ percentile of the within-burst envelope); and burst duration (time from onset to offset) are also reported.

In order to explore the dynamics that occur around burst initiation, maintenance, and termination we performed a time-locked analysis of the evolution of the simulated signals. The timing of concurrent bursts across the network were analysed by using the threshold crossing of an amplitude envelope from a given *sensing* population as reference (Cagnan et al., 2019b) by which to temporally align the data.

We analyse both the probability of a coincident burst occurring across the remaining populations, as well as their relative timing (estimated via the median onset/offset time across all bursts) within a window ±150ms either side of the onset of the burst in the sensing population. This can be used to determine a sequence of changes in the beta amplitudes of each population that preclude the onset and termination of a burst in the reference sensing population. Furthermore, we alter the strength of inputs to the STN to determine how connectivity can dictate the temporal ordering of these events. For the set of results presented in figure 6 we use the STN as the population from which to time-lock the analysis. Differences in the timings of burst onsets and offsets with respect to activity in the STN were computed using two sample t-tests and Bonferroni corrected for multiple comparisons.

### Modelling Closed-Loop Stimulation of Motor Cortex or STN

To explore the impact of phase-locked stimulation on beta bursts, we first simulated a dataset (100s in duration) from the fitted model and identified bursts using the procedures described in section “Definition of Burst Events and their Timings”. We then re-simulated the model with an identical noise process and added an input which had a fixed phase relation with respect to beta activity in either the STN or motor cortex. We analysed models using either cortical stimulation/STN sensing; or cortical sensing/STN stimulation. External stimulation was delivered to the stimulated population for a duration of 300 ms at the onset of a burst in the sensing population.

In order to derive the phase-locked stimulation waveform, we used the analytic phase of the beta activity from the sensing population (*ϕ_REF_* (*t*): the argument of the Hilbert Transform of the unstimulated signal from the sensing population). The stimulation was then given by: *Stim* = *A* sin (*ϕ_REF_*(*t*) + Δ*ϕ*) where Δ*ϕ* sets the phase of the stimulation with respect to the beta activity from the sensing population. *A* denotes the stimulation amplitude and was fixed to yield ¼ of the variance of the intrinsic noise the stimulated population received as an input in the fitted model. For models exploring cortical stimulation, phase-locked stimulation was delivered only to the superficial layer of the cortex (Ali et al., 2013).The phase shift, Δ*ϕ* was swept from −π to +π radians in 12 bins. The resulting power spectra for each stimulation regime was analysed and beta power plotted against phase to yield band-limited amplitude response curves (ARCs) in either lower (14-21 Hz) or upper (21-30 Hz) beta bands. Note that this simulation only mimics closed-loop, as we do not use an on-line phase detection algorithm but rather use the phase computed from the unstimulated data. Identical random input was used to ensure replication of the burst timings for the phase-locked stimulation condition.

In a final set of simulations, we investigate how the ARCs pertaining to the phase-locked stimulation of the motor cortex relative to STN are influenced by the connectivity state of the circuit. To do this we perform the same simulation and analysis as above but modulate the strength of connectivity of the hyperdirect and pallido-subthalamic pathways. The percentage change in amplitude from the median is then used to plot the ARC. We summarise changes to the ARC by reporting the minimum and maximum power, and then plotting these against the strength of connectivity. As above, bursts were predefined according to the unstimulated data.

## Results

### Results of Model Fitting to Electrophysiological Signals in Parkinsonian Rats

Electrophysiological recordings of ECoGs and subcortical LFPs (figure 1A) made in 6-OHDA-lesioned rats were used to constrain a biophysically plausible model of population activity within the Parkinsonian cortico-basal ganglia-thalamic circuit (figure 1B), using a procedure previously reported in West et al. (2019a). The structure of this circuit model, a subset of the data to which it was fit (autospectra only), and the resulting fits are presented in figure 1B. Model parameters were optimized in order to best explain the observed local oscillatory dynamics at each population estimated via their auto-spectra, as well as the interaction between them as measured with directed functional connectivity (estimated using NPD, see methods). For the full set of data features and the resulting fit, see supplementary information I. Properties of the ‘hidden’ unobserved nodes (figure 1B insets indicated with black solid lines) were inferred from the recorded data. Several empirical features of the data were well reproduced by the fitting procedure, such as: 1) widely synchronized beta oscillations across the network (figure 1B; inset); 2) significant feedback of beta oscillations from subcortex to cortex (supplementary information I); 3) bursting behaviour similar to that seen in traces recorded in vivo (figures 1A and C). However, model fits did not well reproduce the higher frequency activity (30-60 Hz) present in the empirical data (figure 1B; inset). The remainder of the results in this paper explore the dynamics of the fitted model, how changes in network state (in terms of the configuration of its constituent connectivity strengths) affect spontaneous activity, and finally how phase-locked stimulation can be utilized to interact with rhythmic activity in the model.

### In-silico Inactivation Experiments: Cortico-Basal Ganglia-Thalamic Connectivity Determines the Power of Subthalamic Beta Rhythms

We performed a set of in-silico inactivation experiments to determine how connectivity within the cortico-basal ganglia-thalamic circuit can lead to emergence of beta rhythms in the STN. Single connections in the model were systematically removed (by setting their weight to zero), and then the resulting changes in beta power in the STN were analysed (figure 2). Three out of ten connections (STR → GPe, GPe → STN, and STN → GPe) acted to substantially promote STN beta oscillations, such that their removal from the model caused > 60% loss in power. Notably, all three connections lay along the indirect pathway and involved the GPe, thereby suggesting that GPe is an important determinant in the promotion of beta oscillations in the STN (figures 2A and B). The cortico-striatal pathway, and the reciprocal connections linking cortex and thalamus, also promoted STN beta, but by considerably less (10% to 30% reduction in power) than those connections targeting or arising from the GPe (figure 2B).

**Figure 2.**
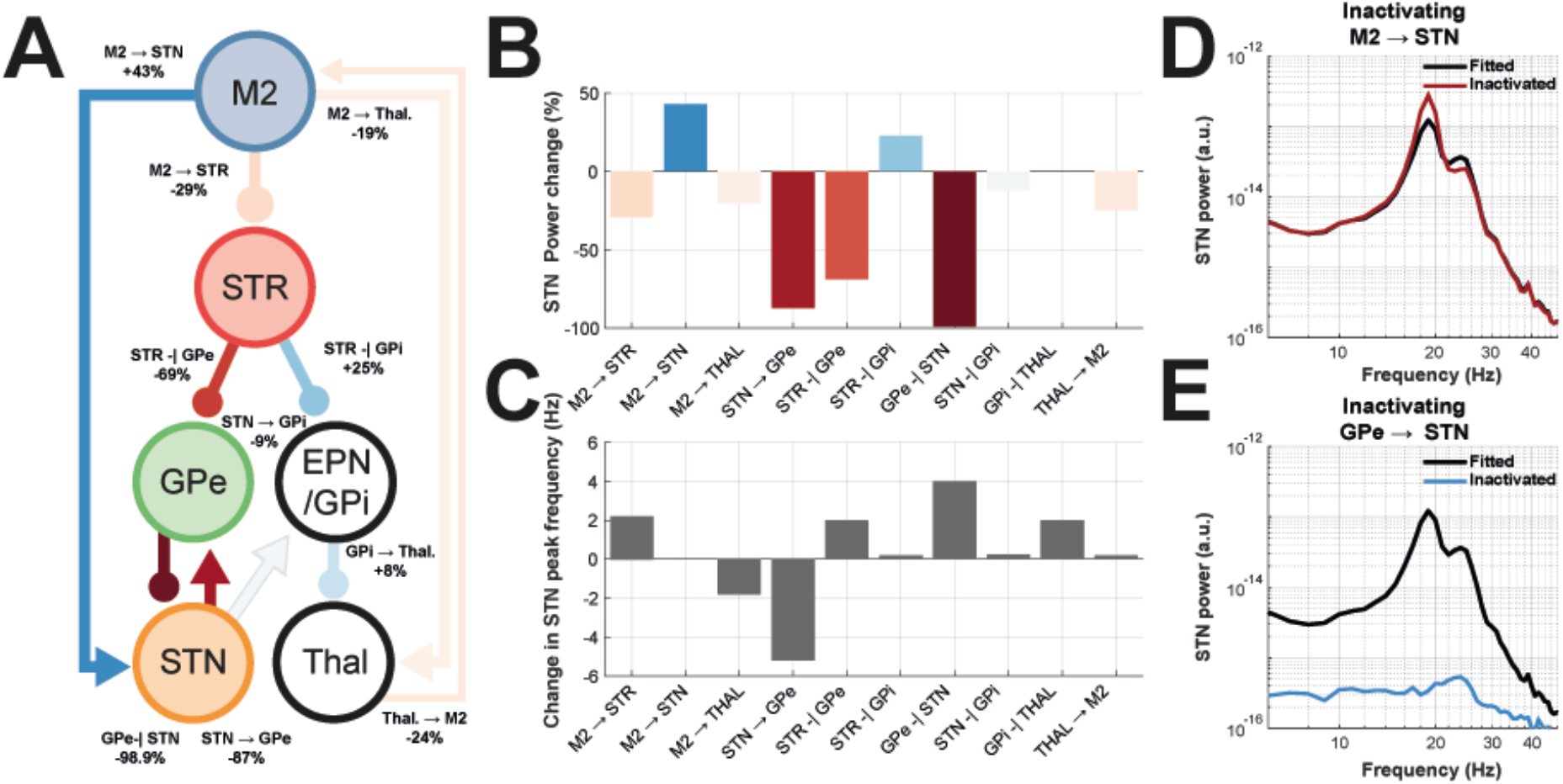
Simulated inactivation experiments to determine the contribution of individual connections in the cortico-basal ganglia-thalamic circuit to the power of STN beta oscillations. Connections in the fitted model were individually removed and the resulting change in simulated STN beta power measured (simulation duration 32 s). **(A)** Schematic of the results of the simulated inactivation experiments. Connections in the circuit are colour coded to indicate the change in 1430 Hz beta power in the STN (red or blue indicating a reduction or increase following lesion, respectively). The actual percentage change is annotated alongside each connection. **(B)** Bar plot of the percentage change in STN beta power from the fitted model following inactivation of each connection. **(C)** Bar plot of the changes in STN peak frequency (within the beta band) from the fitted model following inactivation. **(D)** Power spectra for STN population activity in the fitted (black line) and model with inactivation of hyperdirect pathway (red). **(E)** Same as (D) but for inactivation of the pallido-subthalamic connection (blue line).

In contrast, two connections in the model (M2 → STN and STR → GPi) had suppressive effects on STN beta oscillations, such that their removal resulted in > 25% increases in power (figures 2A and B). Thus, in terms of beta oscillations in STN, the influence of connections involving the hyperdirect and direct pathways appears to be opposite to those involving the indirect pathway. The basal ganglia output pathway (GPi → Thal) was responsible for a weakly suppressive effect (< 10% increase in power; figures 2A and B) and its exact influence is likely determined by the balance of converging inputs from direct/indirect pathways. Using the same set of in-silico inactivations, we also tested for changes in the peak frequency of STN oscillations (figure 2C). Although both STN → GPe and GPe → STN connections promoted STN beta power, they had opposite effects on STN beta frequency: inactivation of STN → GPe decreased frequency by 5 Hz, whereas removal of GPe → STN increased frequency by 4 Hz. In further contrast, removal of the beta suppressing M2 → STN connection did not alter the peak frequency of oscillations in the STN.

In summary, this set of simulations identified that the GPe → STN connection is the primary promoter of STN beta oscillations, whereas the M2 → STN connection acts to suppress STN beta oscillations. Taken together, these data suggest that the influence of the pallido-subthalamic pathway on pathological beta oscillations acts in competition with that of the hyperdirect pathway. The majority of the following results will focus on simulations investigating the effects of modulating pallidal and hyperdirect inputs to the STN.

### Hyperdirect Cortical-Subthalamic Inputs Can Suppress STN Beta Power and Induce a Switch to Higher Frequency Activity

Next, we investigated how gradual changes in the strength of input to the STN from either the cortex or GPe can act to induce transitions in rhythmic behaviour of the model. In figures 3A, B and C, simulated spectra are shown for when the hyperdirect pathway strength (M2 → STN) was altered from the fitted model (connection weight scaled from 10% to 1000% change). As reported in the previous section, weakening of the M2 → STN hyperdirect connection increases the amplitude of subthalamic beta (figure 3A and D). However, strengthening of the same pathway beyond that of the fitted strength (i.e. greater than 100% connection strength) exposes a switching behaviour in the model (figure 3D). This transition involves a switch between lower (14-21 Hz) and higher beta frequencies (21-30 Hz) that occurs at around 160% hyperdirect strength. Beta power in the STN is at a minimum close to this transition in frequency. At connection strengths above 500% of the fitted value, the model is in a hyper-excitable regime reflecting an exponentially divergent state. Simulations that yielded peaks in power that were greater than 200% of that found in the fitted model are treated as outside of the dynamical range of interest and excluded from the remainder of the analyses.

**Figure 3.**
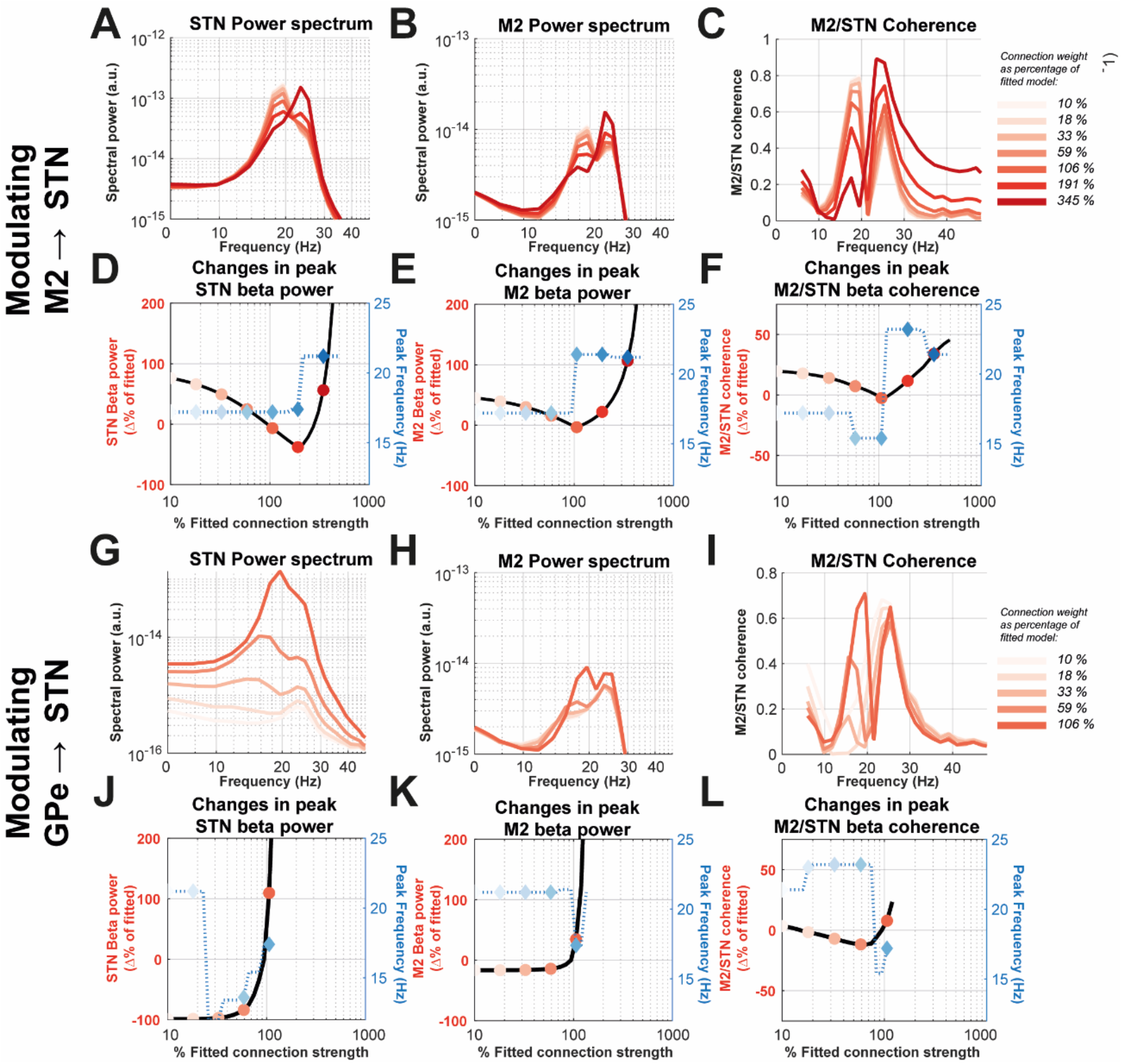
The effects of modulating the strength of STN inputs from the excitatory hyper-direct projection (A-F; M2 → STN) and inhibitory pallido-subthalamic projection (G-L; GPe → STN) upon the power spectra of STN and M2 population activity, and the coherence between them. The strengths of the two connections were varied independently and 256 s of data was simulated from the resulting models. Changes in connection strength are denoted as a percentage of that in the fitted model (i.e. at 100%). Connections were modulated in the range 10% to 1000%. Models yielding an increase in STN beta power greater than 200 % were not analysed. **(A)** The power spectra of the STN resulting from a modulation of the hyperdirect connection. Plots are colour coded according to the key (inset) and matches the markers plot in (D-F). **(B)** Same as (A) but for M2 power spectra. Decreasing the strength of this connection decreases the frequency and amplitude of beta. **(C)** Same as (B) but for STN/M2 magnitude squared coherence. **(D)** Plot of peak beta (14-30 Hz) amplitude (left axis; reds) and peak beta frequency (right axis; blues) in the STN under modulation of hyperdirect input strength. Note X-axis is on a logarithmic scale. **(E)** Same as (D) but for M2 beta power. **(F)** Same as (D) but for STN/M2 coherence. **(G-L)** Plots equivalent to (A-F) but for simulations modulating the pallido-subthalamic connection.

A similar relationship was found when analysing the effects of hyperdirect pathway upon power in the motor cortex where a switch to high beta activity in the motor cortex accompanies an increase in cortical beta power (figure 3B and E). The impact of the hyperdirect pathway upon the magnitude of STN/M2 coherence (figure 3C and F) was also non-monotonic: at strengths close to the fitted value, (i.e. around 100%) coherence is at its minimum. Note that STN beta power approaches its minimal value at stronger connection strengths (~160%). Weakening or strengthening the M2 → STN pathway from this point increases the magnitude of coherence. Attenuating the connection to the smallest value that was simulated (~10% of the fitted strength) increases STN/M2 coherence by approximately 20%. This data can help reconcile the paradoxical findings that dopamine depletion can lead to synaptic weakening of glutamatergic inputs to the STN (Mathai et al., 2015; Chu et al., 2017) whilst at the same time lead to heightened functional connectivity of the same pathway (West et al., 2018).

### Pallidal Inputs to the STN Promote Beta Rhythms

We next modulated the strength of the GPe → STN connection. Simulations presented in figure 3G and J demonstrate the existence of a transition point, close to 32% of the fitted weight, at which there is a shift in beta to higher amplitudes and lower frequencies (transition of peak frequency in the STN from 21 Hz to 12 Hz). Above this point the system is very sensitive to small changes in the connection strength, as STN beta power increases exponentially. Strengthening the inhibition above 125% of the fitted strength results in power exceeding 200% of the fitted, and as above, theses simulations are excluded from the remainder of the analysis. Beta power in the motor cortex (figure 3H and K) yielded a similar response to the STN, with increasing pallidal inhibition inducing a shift to lower frequencies and an exponential increase in power, but with the switch occurring with stronger connection strengths than that required for the STN-closer to the connection strength in the fitted model (~100%).

The effect of strengthening pallido-subthalamic inhibition upon cortico-subthalamic synchronization (figure 3I and L) was found to be biphasic, similar to that found with respect to hyperdirect pathway. Coherence was increased by ~20% when pallidal inhibition was both weakened or strengthened. Again, this switching point was close to the fitted value (90%) at which M2/STN coherence showed a rapid increase and a transition to lower frequencies. The increase in coherence was found to lag that of STN power and exhibited less sensitivity to increases in connectivity.

### Different Feedback Loops Modulate Cortical and Subcortical Beta Activity and their Synchronization

Alterations in the network state likely result from changes in the strength of multiple interacting connections. To explore this hypothesis, we simulated data from 500 models with random strengths drawn from the (inflated) posterior distributions of the fitted model (see methods). We investigate five reciprocal loops shown to be important in maintenance of rhythmic activity (depicted in figure 4A):

1. direct loop: M2 → STR → GPi → Thal. → M2;
2. indirect loop: M2 → STR → GPe → STN → GPi → Thal. → M2;
3. hyperdirect loop: M2 → STN → GPi → Thal. → STN;
4. thalamocortical loop: M2 → Thal. → M2;
5. pallido-subthalamic loop: GPe → STN → GPe.

**Figure 4.**
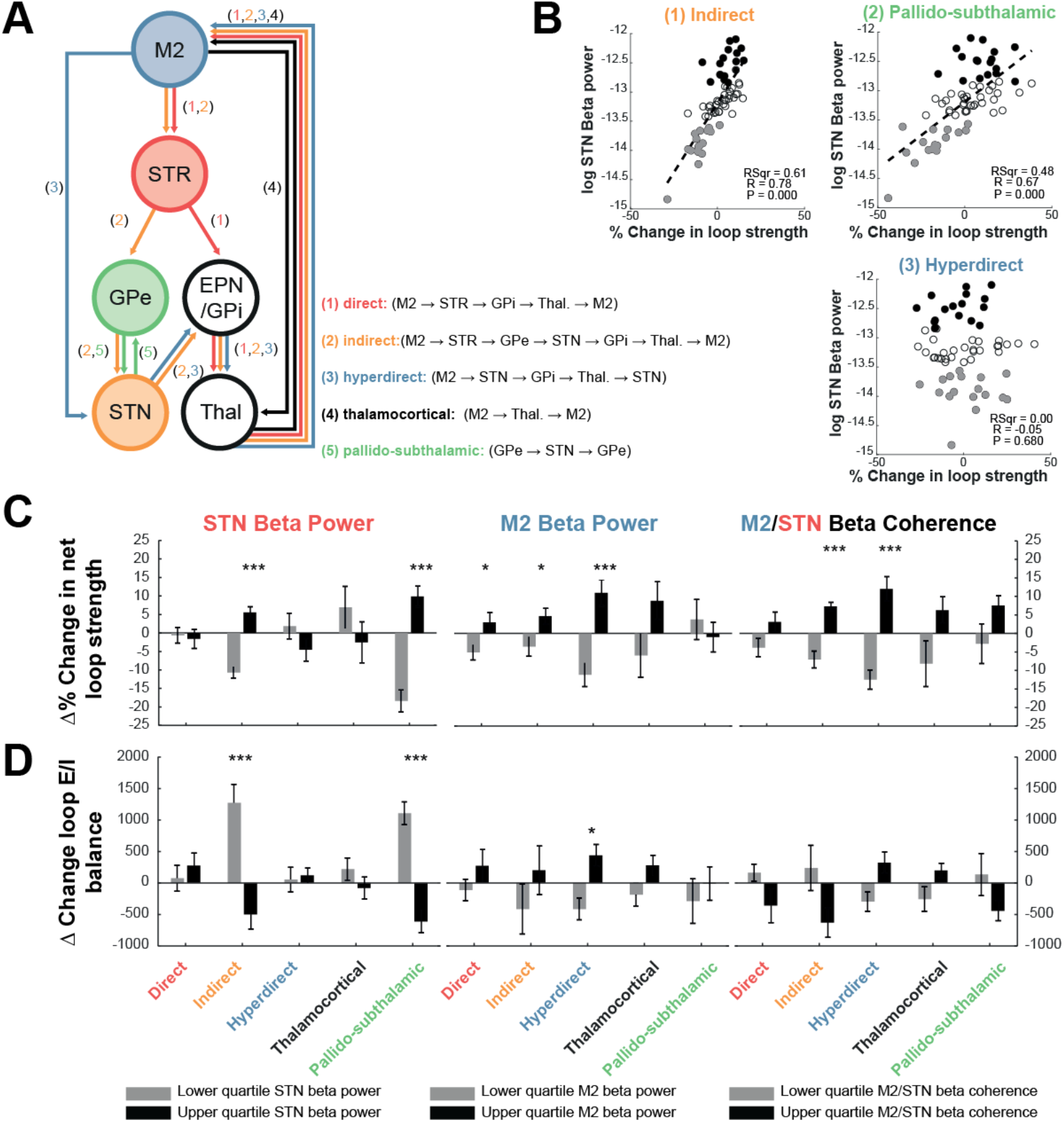
Contributions of loops within the cortico-basal ganglia-thalamic circuit towards the expression of beta (14-30 Hz) power in the STN and motor cortex, as well as their coherence. Random models were generated by drawing 500 samples of connectivity parameters from the (inflated) posterior distributions of the fitted model. These models were then simulated for 30s and the spectral properties of their outputs analysed. **(A)** Circuit diagram indicating the various loops under investigation: direct, indirect, hyperdirect, thalamocortical, and pallido-subthalamic loops. The individual connections contributing to each of the five loops are given by the annotations and colour coding given in the key (inset). **(B)** Scatter plots of *loop strength* vs the resulting STN beta power. Samples in the upper and lower quartiles of beta power are marked in black and grey respectively. Where there was significant correlation (Spearman’s coefficient; Bonferroni corrected α* = α/n; where n = 15 separate tests) a linear regression is plot and resulting statistics inset. This is a selection of graphs- for the full results please see supplementary information II and III. **(C)** Bar graphs of the mean percentage change in *loop strength* associated with models yielding changed in STN beta (14-30 Hz) power, M2 beta power, and STN/M2 coherence below or above the 1^st^ and 4^th^ quartiles (black and grey respectively), and in the first, second, and third columns respectively. Differences in connectivity were tested using student t-tests with Bonferroni corrected significance thresholds (α* = α/n; where n = 30 separate tests; (*: P < 0.05; **: P < 0.01; ***: P < 0.001). Error bars give the standard error of the mean. **(D)** Same as (C) but estimating differences in *excitation/inhibition balance.* Positive values indicate an increase in excitation.

The degree of connectivity within these loops is summarised using either the *net loop strength* (sum of the absolute connectivity strengths) or *net loop E/I balance* (sum of the signed connection strengths; see methods). The differences in these parameters between the models yielding STN beta power, M2 beta power, and STN/M2 coherence below or above the 1^st^ or 4^th^ quartile is presented in figures 4C and 4D.

Results in figure 4C show that for models yielding strengthened STN beta power (1^st^ column) there is an increased loop strength in both indirect loop (−16.3 ±8.8, t(33) = −7.5, P < 0.001, α* = 0.003) and the pallido-subthalamic loop (−28.3 ±16.7, t(33) = −6.9, P < 0.001, α* = 0.003). These changes are associated with an overall increase in inhibition of the two pathways (figure 4D). Correlation analysis of both pathways (figure 4B; supplementary information II and III) confirms the existence of a strong linear relationship between the strengths of these pathways and the spectral features in question. Importantly, M2 beta power is sensitive to different pathways (figure 4C; 2^nd^ column), with increased loop strength of the hyperdirect loop yielding the most prominent change (−22.1 ±19.3, t(33) = −4.7, P = 0.001, α* = 0.003). This change was associated with an overall increase in loop excitation (−8.5 ±10.0, t(33) = −3.5, P < 0.05, α* = 0.003) and in agreement with the results reported in the previous sections. Analysis of models with increased M2/STN coherences demonstrated an enhancement of the hyperdirect pathway (−24.5 ±17.1, t(33) = −5.8, P < 0.001, α* = 0.003) and indirect pathways (−14.3 ±10.4, t(33) = −5.6, P < 0.001, α* = 0.003) which again was associated with an increased excitatory drive in these pathways.

### Pathway Strength Dictates the Probability of Burst Coincidence Across the Network

We next investigated how inputs to the STN alter the evolution of transient beta activity across the network (Tinkhauser et al., 2017b; Cagnan et al., 2019b). To this end, we first estimated the probability of detecting coincident bursts in two populations i.e. given that a burst has been sensed in one population, what is the probability that a coincident burst can be observed in another population? (figure 5A). In general, bursts are highly synchronous across the network with the majority of analyses showing that, in the fitted model, there is over a 50% chance of bursts coinciding within ±150 ms of the onset time at the sensing population. Importantly, these results show that the choice of site at which the reference burst is defined is important in determining the probability of whether a concurrent burst will be observed at another site.

**Figure 5.**
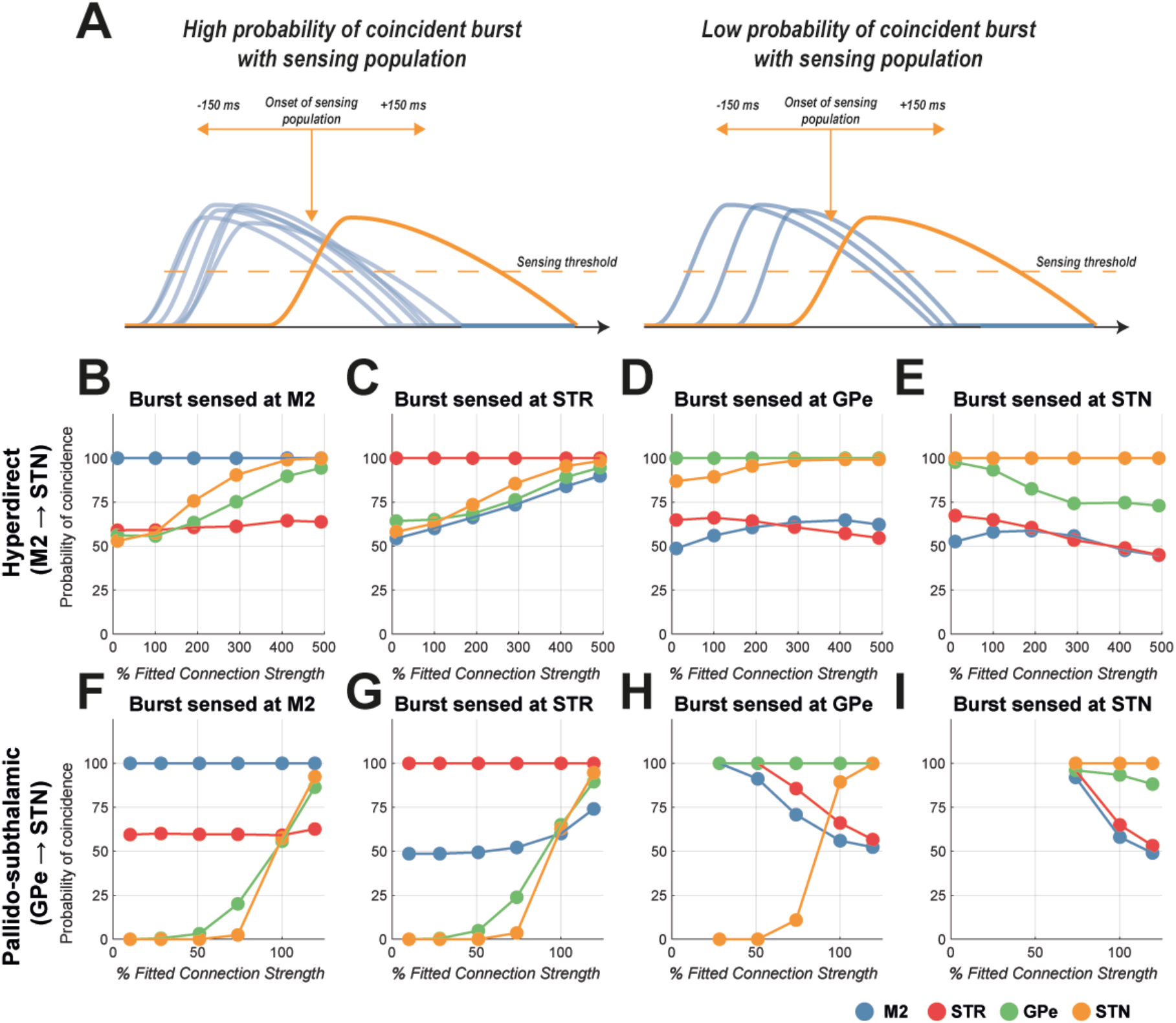
Burst coincidence when beta bursts from different populations are used to define burst onset and the analysis window. The strength of the hyperdirect and pallido-subthalamic projections were modulated from a minimum of 10% of the fitted parameter, up to a maximum that was found to evoke 200% beta power in the STN. **(A)** The probability of coinciding bursts (blue; identified according to the criteria set out in the methods) was calculated within a window 150 ms ± the onset time of a burst detected in the sensing population (yellow). Note that the sensing site will, by definition, have a 100% probability of burst occurrence. **(B, C, D and E)** Analysis of how the probability of burst coincidence changes as a function of hyperdirect pathway strength, when using activity in the M2 (blue), STR (red), GPe (green), or STN (yellow) respectively as a sensing site for detection of beta bursts. **(E, F, G and H)** Same as (A-D) but for modulation of the pallido-subthalamic connection. The equivalent changes in burst properties can be seen in supplementary information IV.

When strengthening the hyperdirect pathway it was found that there was a high probability of bursts across the network coinciding with those detected in the cortex, with > 95% of bursts detected in M2 coinciding with bursts in the STN or GPe (figure 5B, at 500% connection strength). However, if bursts are instead sensed at the STN (figure 5E), increasing hyperdirect strength has the opposite effect: reducing the probability of a coincident bursts being detected at M2, STR, or GPe to below 75% (at 500% connection strength). This result suggests that in the presence of strong cortical inputs to the STN, the majority of suprathreshold beta activity is propagated to STN or GPe, but also that lower amplitude activity (i.e. that are not detectable as a burst in the cortex) becomes sufficient to trigger a high amplitude burst at the STN.

Next, we did the same analysis but when modulating the pallidal-subthalamic pathway (figure 5F). This showed that if this pathway is weak (i.e. less than 50% strength of that in the fitted model), there was a less than 5% probability that bursts sensed at M2 were coincident with those detected in either the STN or GPe (figure 5F, at 10% connection strength). However, when pallidal inputs to the STN were strengthened (and the overall STN beta power was amplified), there is an increased probability of bursts across the network being synchronous with those detected at M2 (greater than 80% probability at 125% pathway strength; figure 5F). If the sensing site is switched to the STN (figure 5I), then again results show an opposite effect to that observed in M2: when the pathway is at its weakest (and still sufficient to generate detectable bursts), then greater than 95% of STN bursts coincide with activity detected in the M2, STR, and GPe. Interestingly, strengthening this input further acts to decrease the probability that bursts sensed in the GPe or STN are coincident with activity in the cortex or striatum (figures 5H and I, less than 75% coincidence at connections greater than the fitted strength). These results suggest that increased pallidal-subthalamic inhibition can permit beta bursts in the STN or GPe to be maintained in the absence of large suprathreshold activity in the motor cortex.

### The Timing of Beta Burst Propagation Across the Network is Altered by Strength of Inputs to the STN

Having identified how bursts coincide across the network, we next analysed the relative timing of burst activity in M2, STR, GPe, and STN and how they change with altered input to the STN. The timings of the burst onsets and offsets (depicted in figure 6A) are given with respect to the STN beta burst onset (i.e. at t = 0). In figure 6B, an example set of beta bursts (defined with a narrowband filter at 18 ±7.5 Hz), simulated from the fitted model, show a cascade of activity with timings that are summarised in figure 6C.

**Figure 6.**
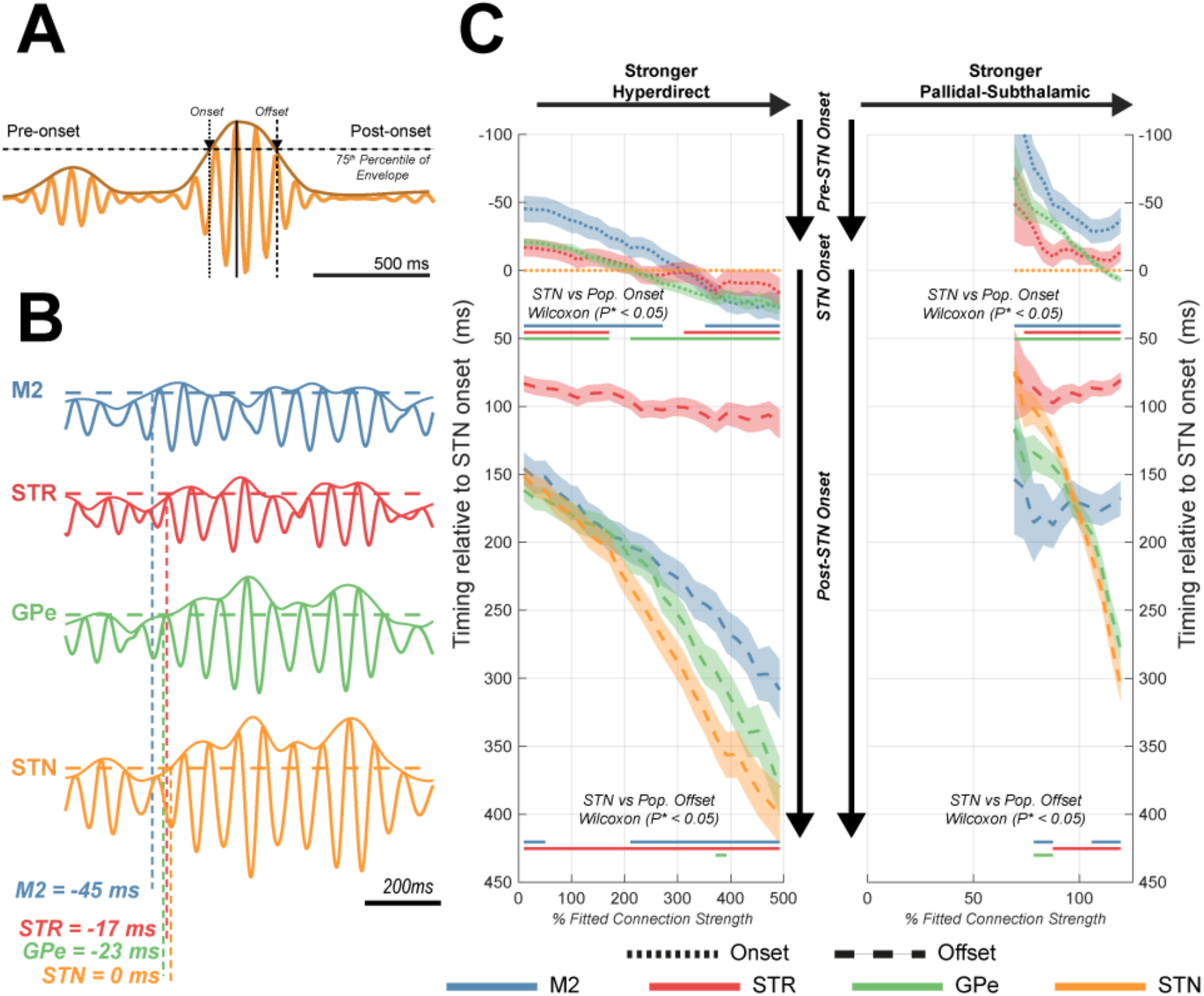
Time-locked analysis of simulated beta bursts across the cortico-basal ganglia-thalamic circuit following modulation of STN inputs via excitatory hyperdirect (M2 → STN) projections and pallido-subthalamic inhibition (GPe → STN). **(A)** Periods of high amplitude beta activity (burst events) were identified by thresholding the band-filtered envelope at the 75^th^ percentile of the data (per population and identified in the simulations of the fitted model). Bursts timings were estimated by the threshold crossing onset and offset. **(B)** Example segment of simulated data shows the propagation of burst activity across the network, with the threshold crossings (i.e. burst onset) of each population in the example annotated. All timings are set relative to the STN burst onset at t = 0 ms. **(C)** Ribbon diagram of the changes in burst timings following modulation of the hyperdirect (left panel) and pallido-subthalamic (right panel) connections. Colour coded lines for each node in the network are given to show the median burst onset (dotted) and offset (dashed) times with borders indicating the 95% confidence interval. The strength of either pathways was modulated from a minimum of 10% of the fitted parameter, up to a maximum that evoked < 200% beta power in the STN. Bold lines indicate simulations in which M2, STR, or GPe bursts onsets/offsets were significantly different from that measured in the STN (t-test, Bonferroni corrected α* = α/n; where n = 18 separate tests).

Figure 6C shows that the timing of the beta bursts across the motor circuit (relative to the onset at the STN) is altered depending upon the strength of input to the STN from cortex or GPe. In the fitted model (i.e. at 100% connection strength), burst onsets occur sequentially: beginning with median onset times at M2 at −36 ms, GPe at −19 ms, STR at −11 ms, prior to STN onset at 0 ms. Bursts began terminating at STR at 92 ms, followed by a gap of ~80 ms (equivalent to ~1.5 cycles of beta), and then a rapid termination of the burst in the remainder of the circuit (M2 at 171 ms, GPe at 179 ms, and STN at 180 ms). At all connection strengths that were investigated, striatal bursts terminate from 50ms to 100ms prior to those in the remainder of the circuit suggesting that they may not be required for the maintenance of STN beta activity.

The effect of the hyperdirect M2 → STN connection on burst timing is shown in figure 6C (left panel). This analysis shows that the strength of this pathway determines whether cortical burst onsets precede or succeed those in the STN (*M2 onset times min. vs max. hyperdirect connection strength:* 73 ±14.5 ms, t(573) = 9.84, P < 0.001), with the strongest hyperdirect connection postponing M2 onset to +27 ms. These effects are accompanied by a delaying of bursts offsets (*STN offset times for min. vs max. hyperdirect connection strength:* 249 ±19.5 ms, t(1029) = 25.1, P < 0.001).

Similarly, the effects of the pallido-subthalamic GPe → STN connection are shown in the right panel of figure 6C. As might be expected, increasing this pathway’s strength from the fitted model, brings GPe bursts’ onset closer to that found in the STN (*GPe onset times for fitted. vs max. pallido-subthalamic connection strength:* 22 ±4.0 ms, t(1398) = 11.0, P < 0.001). Additionally, strengthening pallido-subthalamic input to its maximum elongates bursts in the STN by delaying burst termination by an average of 125 ms (equivalent to ~2 cycles of beta) when compared to the fitted model (*STN offset times for fitted vs max pallido-subthalamic strength:* 125.0 ±16.5 ms, t(1523) = 14.8, P < 0.001) with activity that is tightly followed by the GPe. At the same time the quenching of beta activity in the cortex is unchanged (*M2 offset times for fitted vs max pallido-subthalamic strength:* 3.4 ±3.5 ms, t(895) = 0.28, P = 0.78). These changes indicate that increased pallidal inhibition induces a divergence of activities in STN and GPe that extend for one to two cycles beyond the termination of beta bursts in the cortex.

### Phase-Locked Stimulation of Motor Cortex Triggered by Beta Bursts Sensed in the Subthalamic Nucleus

In the next set of simulations, we explored how the timing of cortical activity modulated STN beta bursts by modelling the effects of phase-locked stimulation delivered to the cortex. Beta activity sensed at the STN was used to control phase locked stimulation. Assuming that stimulation is delivered to cortex non-invasively e.g. using transcranial alternating current stimulation, stimulation was modelled as input to the superficial layers of cortex (methods; figure 1B). Figure 7A demonstrates that STN beta activity can be either amplified or suppressed depending on the relative phase of the stimulation signal delivered to M2. When close to anti-phase (i.e. a phase difference of 180° equivalent to when a peak in M2 peak is aligned to a trough in STN), there is a promotion of STN beta amplitude, and a suppression when in-phase (i.e. a phase difference of 0° equivalent to an alignment of M2 and STN peaks). These effects presumably occur due to constructive/destructive interference (Rosenblum and Pikovsky, 2004; Witt et al., 2013), with the precise phase of the effect matched to the transmission delays involved in propagation of the rhythmic activity. Cortical stimulation impacts both the cortical and subthalamic power spectra, as well as the coherence between them (shown in supplementary information V) with a response that is dependent on the precise phase difference with respect to ongoing activity in the STN. Despite the stimulation phase being derived from 18 ± 7.5 Hz STN activity, phase locked stimulation differentially affects spectral peaks at low (14-21 Hz) and high (21-30 Hz) beta frequencies in STN and M2 depending on the stimulation angle.

**Figure 7.**
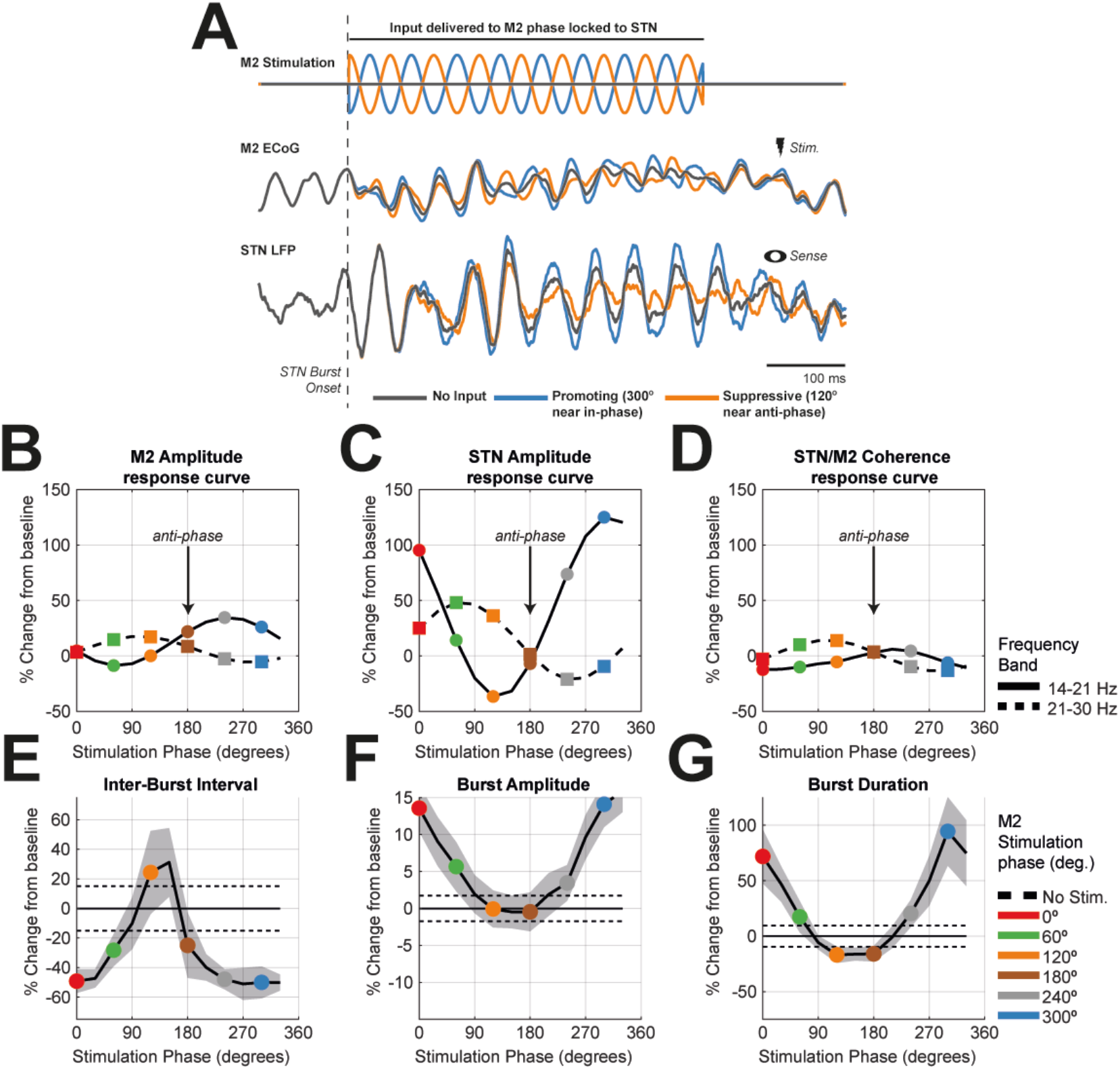
Phase locked stimulation of motor cortex when sensing bursts in the STN of the fitted model. Bursts were defined using the narrowband activity (18 ±7.5 Hz) simulated at the STN. **(A)** Example simulation within a single STN burst: a high amplitude STN burst was detected from simulations of the fitted model (burst onset indicated by grey dashed line and time courses of activity across the network shown by the bold grey line). Phase locked inputs were then added in a second repeat simulation which used identical noise inputs to the network to ensure burst timings were replicated. The timings of burst onsets were used to trigger a 300ms sinusoidal input to cortex (example shown in orange and blue; top) with its phase manipulated to have an offset with respect to that of ongoing beta activity in the STN. We show M2 stimulation phases yielding maximal promotion (blue; close to in-phase with STN-i.e. peak to peak) or suppression (orange; close to anti-phase with STN-i.e. peak to trough) of STN low beta power (14-21 Hz). **(B, C, D)** ARCs corresponding to the changes in spectral power (for corresponding spectra see supplementary information V) indicating the modulation of low beta (14-21 Hz; bold, circles) and high beta (21-30 Hz; dashed, squares) frequencies. Peaks are plot as a percentage difference from the fitted model (i.e. the unstimulated condition). Coloured symbols correspond to the phase bins given in the key (inset). Changes in burst properties (see supplementary information VI for schematic of methods and additional changes to burst properties) are shown for the inter-burst interval **(E)**, burst amplitude **(F)**, and burst duration **(G)**.

These phase specific effects of stimulation can be summarised by constructing ARCs which show that there are differences in the maximally suppressive phases of low beta in the M2 and STN: in M2 this occurs close to a stimulation phase value of 60° (figure 7B, green circle; 9% reduction of power) versus 120° in STN (figure 7C, orange circle; 37% reduction in power). Secondly, there is a differential response for modulation of high and low frequency beta rhythms in the STN: low beta is enhanced at 300° (figure 7C, blue circle; 125% increase in power); whilst at the same phase STN high beta is close to its most suppressed (figure 7C, blue square; 10% reduction in power). The difference in peak modulation for the two bands is separated by 120°. These results indicate that the range of phases in which it is possible to achieve a suppressive effect for STN low beta (4/12 phase bins) is smaller than that found to be amplifying (8/12 of phase bins). Coherence of the STN/M2 beta shows only a weak capacity to be modulated by phase locked stimulation of M2 (figure 7D; maximum modulation of ±14% from baseline).

### Phase-Locked Stimulation of Cortex can Modulate Properties of Burst Activity in the Subthalamic Nucleus

We next demonstrate how the effects of phase locked stimulation of the cortex upon STN beta power can be explained in terms of changes to the temporal patterning of beta activity (figure 7E, F, and G). The properties of these bursts are depicted schematically in supplementary information VI. We show that stimulation of the motor cortex at phase values which suppress STN low beta, this effect is predominantly related to an increase in the average interburst interval (figure 7F; *maximum suppressive stim phase. vs base.;* −0.1 ±0.24 s, t(265) = −2.86, P < 0.01, α* = 0.01) and the abridgment of burst durations (figure 7G; *maximum suppressive stim phase. vs base.;* 44 ±20.8 ms, t(269) = 4.19, P < 0.001, α* = 0.01), rather than through reduction in burst amplitude (figure 7F; *maximum suppressive stim. phase vs base.;* 6.7 x 10^−8^ ± 5.6 x 10^−8^, t(269) = 2.37, P = 0.0184, α* = 0.01).

### Phase Locked Stimulation of STN Triggered by Beta Bursts in the Motor Cortex

The results presented in the section *Pathway Strength Dictates the Probability of Burst Coincidence Across the Network* suggests that, dependent upon the connectivity state of the network, there can be a high coincidence between cortical and STN beta bursts. We next explored the impacts of switching sensing and stimulation sites such that stimulation was delivered to the STN with respect to the phase of cortical beta bursts. Phase-locked stimulation applied to the STN yields smaller effects upon the power spectra (figure 8A, B, and C). The response to stimulation is largest in the STN and mainly restricted to low beta band activity (figure 8B and E, bold line; maximum of 59% increase in power for low beta vs maximum of 13% increase for high beta). The suppressive effect of low beta was slightly weaker than that seen during M2 stimulation (−31% versus −37% for STN stimulation). Modulation of cortico-subthalamic coherence by STN stimulation was also smaller than that during cortical stimulation, with a modulation of only ±7% from the baseline (figure 8C and F) compared to ±14% for M2 stimulation. Overall, results suggest that STN stimulation may provide more specific targeting of STN low beta activity but lacks the larger effect sizes that M2 stimulation can elicit.

**Figure 8.**
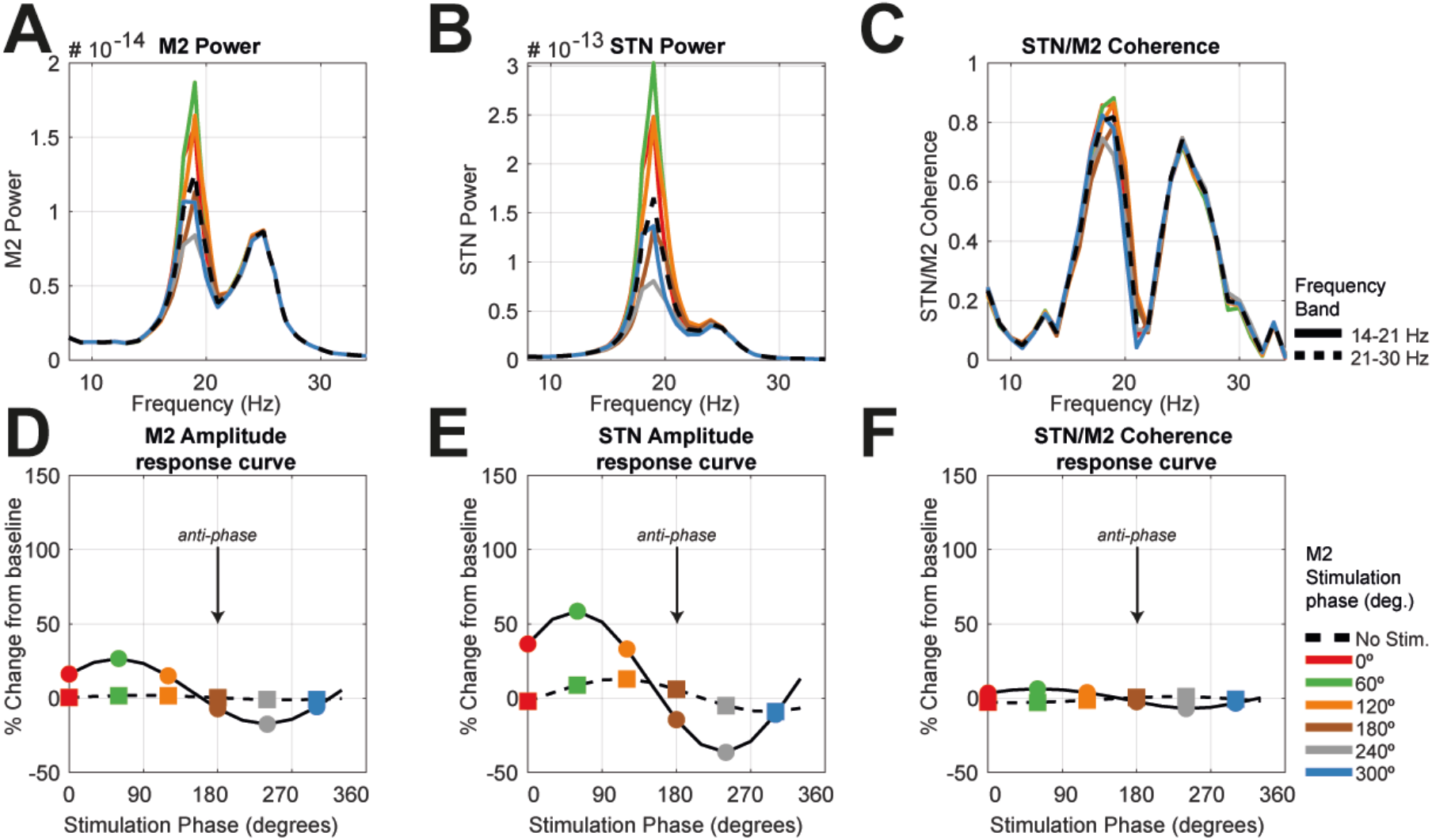
Simulating phase-locked stimulation of STN when sensing bursts in the motor cortex of the fitted model. **(A, B, and C)** Plots of M2 power spectra, STN power spectra, and STN/M2 coherence for six selected phase bins (given in legend). The dashed line gives the spectra from the unstimulated condition (i.e. fitted model). **(D, E, F)** Response curves corresponding to the spectra in the row above indicating the modulation of low beta (14-21 Hz; bold, circles) and high beta (21-30 Hz; dashed, squares) frequencies. Peaks are plot as a percentage difference from the fitted model. Coloured symbols correspond to the sample phase bins of the spectra shown in the above row.

### Modulation of STN Beta Amplitude by Cortico-Subthalamic Relative Phase is State Dependent

In figure 9, we return to the more efficacious stimulation of the cortex and investigate the effect of modulating the strength of inputs to the STN. These analyses show that the profiles of the ARCs are significantly altered by changes in the strength of cortical or pallidal inputs to the STN (examples shown in figure 9A and D respectively). These changes involve both alterations in the maximum suppression and amplification of beta rhythms achieved by stimulation, as well as shifts in the effective stimulation angle. The dependency of ARC properties upon hyperdirect M2 → STN connectivity is shown in figure 9B. Most ARCs indicate an amplifying and suppressive regime, although amplification is always larger than the corresponding suppression (e.g. 149% vs −52% in the fitted model). Analysis of the maximal suppressive/amplifying phase (figure 9C) indicates a shift in the maximally suppressive angle around 160% connection strength, the same transition point that was found to result in a shift to higher frequency beta activity in figure 3. This result likely arises from a shift in pathways responsible for the maintenance of beta oscillations.

**Figure 9.**
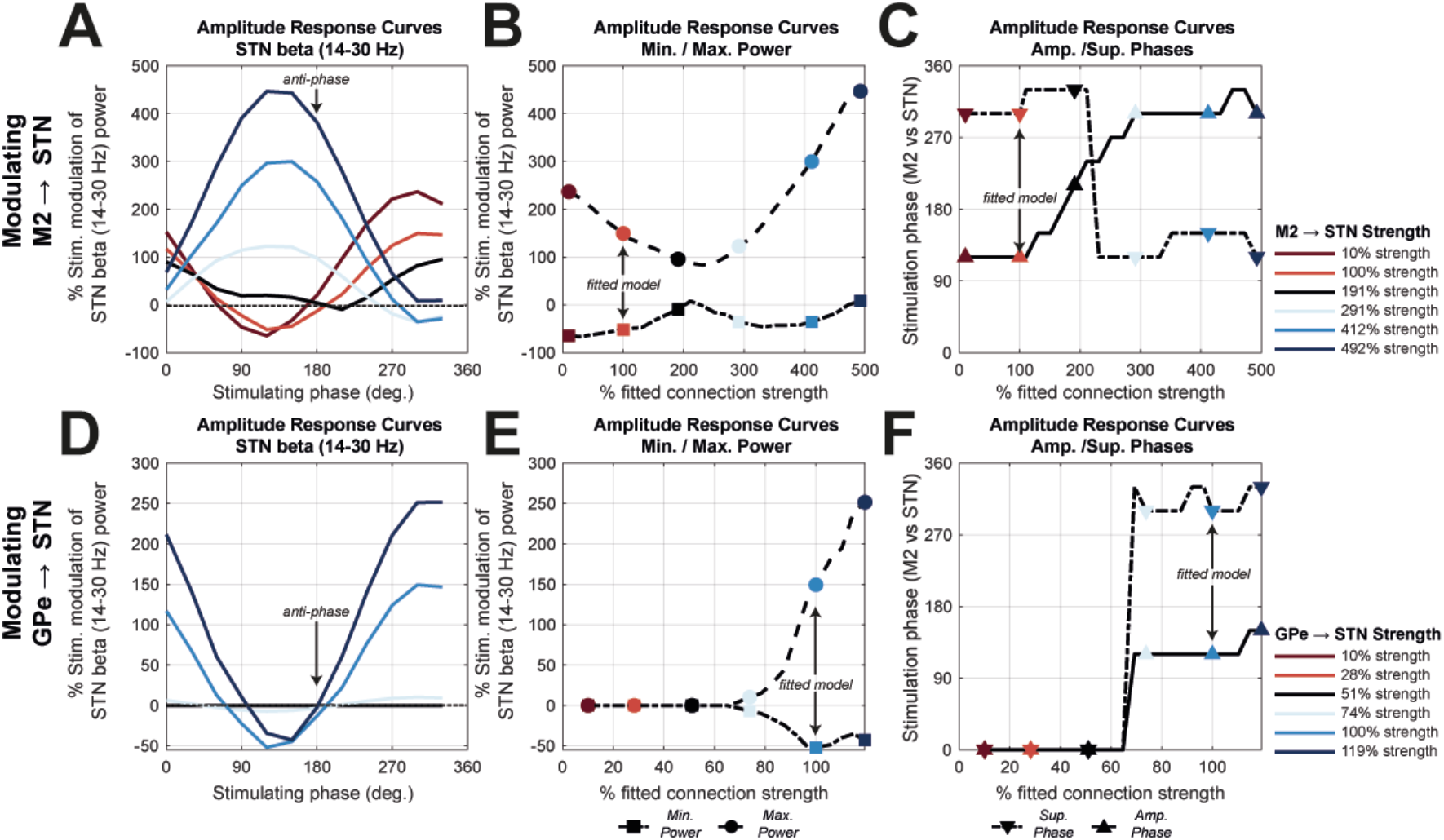
Simulating state dependent changes in the efficacy of phase locked stimulation of motor cortex (M2) when modulating pallido-subthalamic (GPe → STN) and hyperdirect (M2 → STN) connection strength. Connection strengths were varied from 10% of the fitted model up to a maximum that elicited < 200% change in STN beta power. Changes in power are scaled as the percentage difference from the unstimulated for each connection strength. **(A)** Simulated ARCs when the strength of the M2 **→** STN connection was modulated. These ARCs exhibit changes in maximum suppression/amplification, as well as shifts in phase. **(B)** The ARCs were characterised by taking the minimum (dotted line; circles) and maximum (dashed line; squares) power and then plotting these against the corresponding connection strength. The coloured markers correspond to the selected ARCs shown in (A). **(C)** Plot of the maximally suppressive (dashed; downwards arrows) and promoting (bold; upwards arrows) phase versus connection strength. There is a clear transition in phase close to 200% change in connection strength. **(D, E, F)** Same as (A, B, C) but for simulations modulating the GPe → STN connection.

Modulation of the pallido-subthalamic GPe → STN connectivity shows that the response is mostly amplifying, with amplification found to be much larger than suppression in all cases tested (figure 9D and E). Both suppressing and promoting stimulation phases were found to be stable for states associated with significant beta power (i.e. at connection strengths greater than ~60%; figure 9F).

## Discussion

### Summary of Findings

Using a set of in silico analyses, we have shown that the emergence of subthalamic and cortical beta oscillations is under the influence of an interplay of overlapping loops across the cortico-basal ganglia-thalamic circuit. Due to their prominent effect on subthalamic beta power, we focused on inhibitory and excitatory inputs to the STN, arriving from the external pallidum and motor cortex respectively, and used variations in these connections to shift the cortico-basal ganglia-thalamic circuit into different network states. We have shown that these connections also influence the rhythm of subthalamic beta oscillations; with hyperdirect pathway input biasing these oscillations to higher frequencies. We have characterised the intermittent nature of beta oscillations and determined burst coincidence, onset and termination across the motor circuit. This analysis has demonstrated that subthalamic and pallidal beta bursts significantly overlap and given sufficiently strong pallidal inhibition of the STN, will dissociate from those observed in the cortex and striatum. Finally, we have shown that phase-locked stimulation can selectively enhance or suppress beta oscillations depending on stimulation timing, and yields an effect that is strongly dependent on network state (in the context of the balance of excitatory and inhibitory inputs to the STN). Phase-locked stimulation induced an increase in the average inter-burst interval and reduced mean burst durations when subthalamic beta oscillations were suppressed. Extensive effort is currently being devoted to the development of novel stimulation based therapies (Cagnan et al., 2019a). Our results provide valuable insights that can guide the design of neuromodulation strategies that are reactive to changes in brain state.

### The Amplitude of STN Beta Rhythms is Influenced by Alterations in Network Connectivity

In this work, we have shown how subthalamic beta power, a key biomarker used for aDBS (Little et al., 2013; Rosa et al., 2015), can be modulated by shifts in the engagement of different pathways across the motor circuit (figures 2-4). The STN/GPe loop has been previously implicated in basal ganglia pathophysiology and is associated with aberrant STN beta power (Mallet et al., 2008a; Cruz et al., 2011; Tachibana et al., 2011). In our study, we observed that reducing the strength of the pallido-subthalamic connection diminished beta power in the STN, in good agreement with the recent empirical demonstration that suppressing GPe neuron activity decreases the engagement of STN neurons in Parkinsonian beta oscillations (Crompe et al., 2020). In contrast to the influences of STN/GPe connections, we observed that reducing the strength of the cortico-subthalamic hyperdirect connection augmented beta power in STN. Thus, moderate increases in hyperdirect pathway transmission might decrease STN beta, with attendant behavioural benefits, a prediction supported by small-scale studies in Parkinsonian mice (Sanders and Jaeger, 2016). Taken together, the results presented here indicate that the propensity of the STN/GPe subcircuit to enter a resonant state arises from an interplay of pallidal and hyperdirect pathway inputs impinging upon the STN. This matches well with previous computational models (Pavlides et al., 2015; Fountas and Shanahan, 2017) that propose that the STN/GPe loop selectively resonates (Hahn et al., 2014) at beta frequency as a result of inputs arriving from the motor cortex, as well as experimental findings that Parkinsonism predisposes the circuit to synchronization by the cortex (Baaske et al., 2019).

The results of our model reconcile the apparently paradoxical findings that dopamine depletion can result in enhanced cortical-subthalamic coherence (Sharott et al., 2005b; West et al., 2018; Baaske et al., 2019) following synaptic weakening of glutamatergic inputs to the STN (Mathai et al., 2015; Chu et al., 2017). Our work suggests that this effect results from the loss of competing input to STN that acts to increase patterning of the STN by beta activity propagating via the cortico-striatal indirect route. Critically, indirect and hyperdirect loops may leave distinct spectral signatures in terms of the expression of low and high frequency beta, respectively (figure 3).

In contrast to previous work (van Albada et al., 2009; Brazhnik et al., 2016; Reis et al., 2019) we find little evidence to support the significance of the corticothalamic relay (acting in insolation from basal ganglia output) in the maintenance STN beta rhythms (figures 2 and 4). It is however important to note that our study did not use thalamic recordings to constrain model parameters, which may explain its relatively unimportant role in the presented analyses. By expanding on the principle of competing loops (Leblois et al., 2006) to include those formed by the subthalamo-pallidal feedback, indirect, and thalamocortical relay in addition to the hyperdirect and direct loops, we show that maintenance of STN and cortical beta involve distinct loops (figure 4).

### Disrupting Antecedent Cortical Activity May Provide Selective Targeting of Pathological Activity

Beta activity evolves as it propagates through the different loops of the circuit. Beta bursts in the basal ganglia tended to be preceded by bursts in the cortex (figures 5 and 6), a finding in agreement with experimental evidence demonstrating that cortical beta rhythms on average drive those in the STN (Williams et al., 2002; Fogelson et al., 2006; Lalo et al., 2008; Litvak et al., 2011; Sharott et al., 2018). In this study, noise was delivered across subcortical populations and cortex. Therefore, temporal evolution of beta bursts is not explicitly built into the model. Analysis of the timing of burst termination in the model demonstrates how increased pallidal inhibition of the STN can result in burst elongation (figure 6C), offering a candidate mechanism for generation of long bursts that have been correlated with the Parkinsonian state (Deffains et al. 2018; Tinkhauser et al. 2018). A synthesis of the two above findings suggest that long duration beta bursts may begin as either subthreshold or short duration beta bursts in the motor cortex that is later prolonged within the STN/GPe loop (Cagnan et al., 2019b). The circuit will resonate at beta frequencies given sufficient strength of the GPe/STN feedback and a suitable phase alignment with cortex. Intriguingly, beta bursts in this network state tend to be at low beta frequencies (14-21 Hz; figure 3). When the STN/GPe coupling is very high, beta activity can be sustained to such a degree that it may terminate several cycles later than cortical beta bursts (figure 6). Our results give rise to an experimentally testable hypothesis and suggest that beta bursts in the cortex and STN should diverge in duration with increased activity of neurons in the GPe. Critically, interruption of precursor activity in the cortex may be used to quench the states that promote the emergence of heightened STN beta resonance using existing surgical techniques (Swann et al., 2017).

### The Responses of STN Beta Rhythm to Phase Locked Stimulation are State Dependent

At its most basic, therapeutic brain stimulation, such as cDBS for PD, provide stimulation using fixed parameters (i.e. frequency and amplitude) that are chosen to provide clinically significant reductions in patients’ symptoms whilst minimising undesired adverse effects (Castrioto et al., 2014; Cagnan et al., 2019a). More recently, approaches such as aDBS have been proposed to gate the delivery of stimulation to biomarkers associated with a pathological or functional state (Little et al., 2013; Rosa et al., 2015; Swann et al., 2018; Bouthour et al., 2019). This approach has been shown to be effective in reducing PD motor symptoms (Little et al., 2016a) together with improving stimulation side-effects such as speech intelligibility (Little et al., 2016b). However, beta rhythms which are commonly used to control the delivery of stimulation, and also correlate with physiological processes across the cortico-basal ganglia circuit (Khanna and Carmena, 2015; Mirzaei et al., 2017; Shin et al., 2017; Hannah et al., 2019), suggesting that stimulation specificity may need to be further improved.

Phase-locked DBS could provide such a refinement in stimulation specificity by delivering precisely timed stimulation to modulate rhythmic activity together with interregional synchronization. This approach follows on from the fact that neuronal ensembles demonstrate phase locked firing to population activity. Therefore, precisely timed stimulation relative to the rhythmic control signal may act to either enhance or reduce the amplitude of these rhythms depending on stimulation timing (Rosenblum and Pikovsky, 2004; Witt et al., 2013; Holt and Netoff, 2014; Wilson and Moehlis, 2015). Phase locked stimulation approaches have been demonstrated to be effective in modulating tremor (Brittain et al., 2013; Cagnan et al., 2017), supressing beta oscillations in Parkinsonian patients (Holt et al., 2019), as well as in healthy (Peles et al., 2020) and Parkinsonian primates (Sanabria et al., 2020). The parameters required for optimal therapeutic neuromodulation are likely to be brain state dependent (Bergmann et al., 2016; Karabanov et al., 2016; Kahan et al., 2019). In this work we have demonstrated that the optimal stimulation phase and its ability to modulate beta rhythms, is influenced by the strength of synaptic inputs to the STN. This dependency arises from the impact these projections have on synchronous neural activity across the motor circuit (figure 3; Tass, 2000; Weerasinghe et al., 2019). These results suggest that static approaches to selecting stimulation control parameters are likely to be non-optimal in the face of spontaneous shifts in brain state that, for instance, accompany movement or action selection. Future stimulation strategies that can provide dynamic parameterization of stimulation may adopt state estimation techniques (Baker et al., 2014) or dual control algorithms that can provide behaviour responsive delivery of neuromodulation (Grado et al., 2018).

### Targeting Interareal Synchronization with Phase Locked Stimulation

Our simulations demonstrate that when phase-locked stimulation supressed STN rhythms in low beta frequencies, there was a simultaneous enhancement of beta activity at higher frequencies (> 21 Hz) (figure 7), comparable to spontaneous shifts we observed due to increases in the excitatory inputs to the STN via the hyperdirect pathway (figure 3). Similar results have been demonstrated experimentally (Sanabria et al., 2020) where a higher frequency beta rhythm (> 18 Hz) emerged during phasic DBS targeting 11-17 Hz rhythms. These results suggest that phasic suppression of neural rhythms at a certain node may shift circuit activity and promote communication via specific loops such as the hyperdirect pathway. This phenomenon is not limited to the cortico-basal ganglia-thalamic circuit and was also observed in essential tremor patients during phase-locked thalamic DBS (Cagnan et al., 2017). The potential to reshape activity in these circuits is likely to have important implications for motor execution and planning (Frank et al., 2007; Jahfari et al., 2011). These effects can be controlled by careful determination of both the “sensing” site at which pathological activity is detected, as well as the site at which stimuli are delivered (figures 7 and 8). Alternating the “sensing” and “stimulating” nodes between the STN and cortex impacted both the emergence of different rhythms during stimulation and the stimulation effect size: with phase-locked subthalamic stimulation with respect to cortical beta rhythms giving rise to smaller effect sizes than phase-locked cortical stimulation with respect to subthalamic beta. Critically, phase-locked stimulation induced an increase in the inter-burst interval and reduced mean burst durations while having minimal impact on burst amplitude.

### Limitations

All computational models are limited to the range of neuronal features they can describe by the form of the equations used to explain them. In this work we utilize so called “lumped parameter” models that simplify the description of neuronal dynamics by effectively averaging over a very large number of states and parameters. In making this simplification, these models lack the ability to describe many phenomena such as the complex neuronal spiking of basal-ganglia neurons, or their nonlinear integration of inputs (Farries et al., 2010; Amadeus Steiner et al., 2019). The low dimensionality of these models also effectively restricts their dynamic range, with population time constants restricting investigation of the interaction between very fast inputs and slower neuronal responses, such as that imposed by high frequency DBS. Moreover, this model does not capture several rhythms such as higher frequency gamma rhythms (Swann et al., 2018) thought to be important in movement, or low frequency alpha/theta which has been implicated in this circuit in dystonia (Wang et al., 2018). To capture these features, extensions of the model would require the inclusion of populations with time constants suited to propagating different rhythms. Finally, this model is the result of fitting to time averaged features of the original data. It remains to be seen as to whether including parameters of the temporal intermittencies to further constrain model parameters can provide better mechanistic models of the cortico-basal ganglia-thalamic system.

### Conclusions

This study builds upon our understanding of the emergence of beta oscillations in the cortical-basal ganglia-thalamic circuit by elucidating the potential mechanisms by which pathological activity may propagate throughout the recurrent circuits. Our results suggest that heightened beta activity in the STN results from the strengthening of the indirect pathway involving GPe, whilst beta activity in the motor cortex is promoted by the hyperdirect pathway and its thalamocortical return. Analysis of the state dependent changes in beta burst timings suggest that strengthening of pallidal input to the STN leads to beta rhythms in the STN/GPe circuit to become progressively less dependent on cortical input. Finally, we show that subcortical beta rhythms can be modulated by precisely timed stimulation of the cortex, with a response that is non-trivially related to network connectivity. These results support novel approaches to neuromodulation that can adapt stimulation in accordance with changes in brain state. The broad involvement of interareal synchronization in both pathology and functioning of the brain (Schnitzler and Gross, 2005; Bressler and Menon, 2010; Thut et al., 2012) makes phase-locked modulation of brain networks of great relevance to circuits beyond the cortical-basal ganglia-thalamic network and could potentially be applied to those involved in memory, sleep, and decision making.

## Acknowledgements

We would like to thank both Prof. Peter Brown and Ms. Carolina Reis for their helpful comments. We thank Dr. N. Mallet for acquiring some of the primary data sets. This work uses a number of toolboxes, generously developed, maintained, and shared by the community (details in supplementary information VII) and to whose authors we are very grateful.

## Supplementary Information

**Supplementary Information I.**
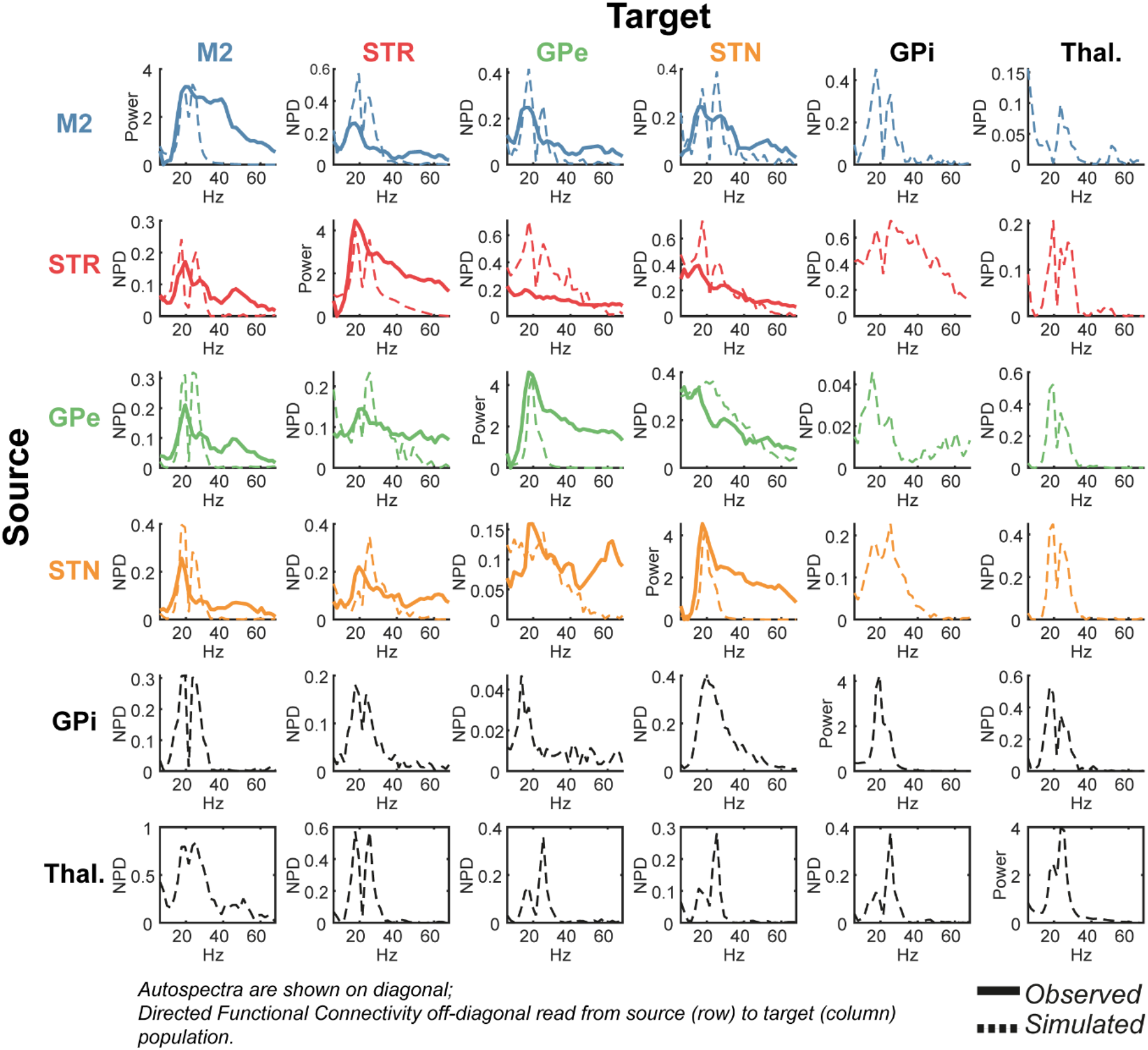
Full set of empirical data features: autospectra (diagonal) and directed functional connectivity (off-diagonal) used in the parameter optimization, and their resulting fits. Empirical data (bold) is taken from a group level analysis of electrophysiological recordings made in the 6-OHDA rodent model of Parkinsonism (Mallet et al., 2008b, 2008a). ECoG was recorded from M2 region of cortex (blue), and LFPs from the STR (red), GPe (green), and STN (yellow). Data is shown prior to Gaussian smoothing that was applied before fitting. Predicted spectra are shown for the hidden nodes at GPi and Thalamus. Autospectra are placed along the diagonal. Directed functional connectivity (Non-parametric directionality; West et al., (2018)) is on the off-diagonal and can be read from source (row) to target (column).

**Supplementary Information II.**
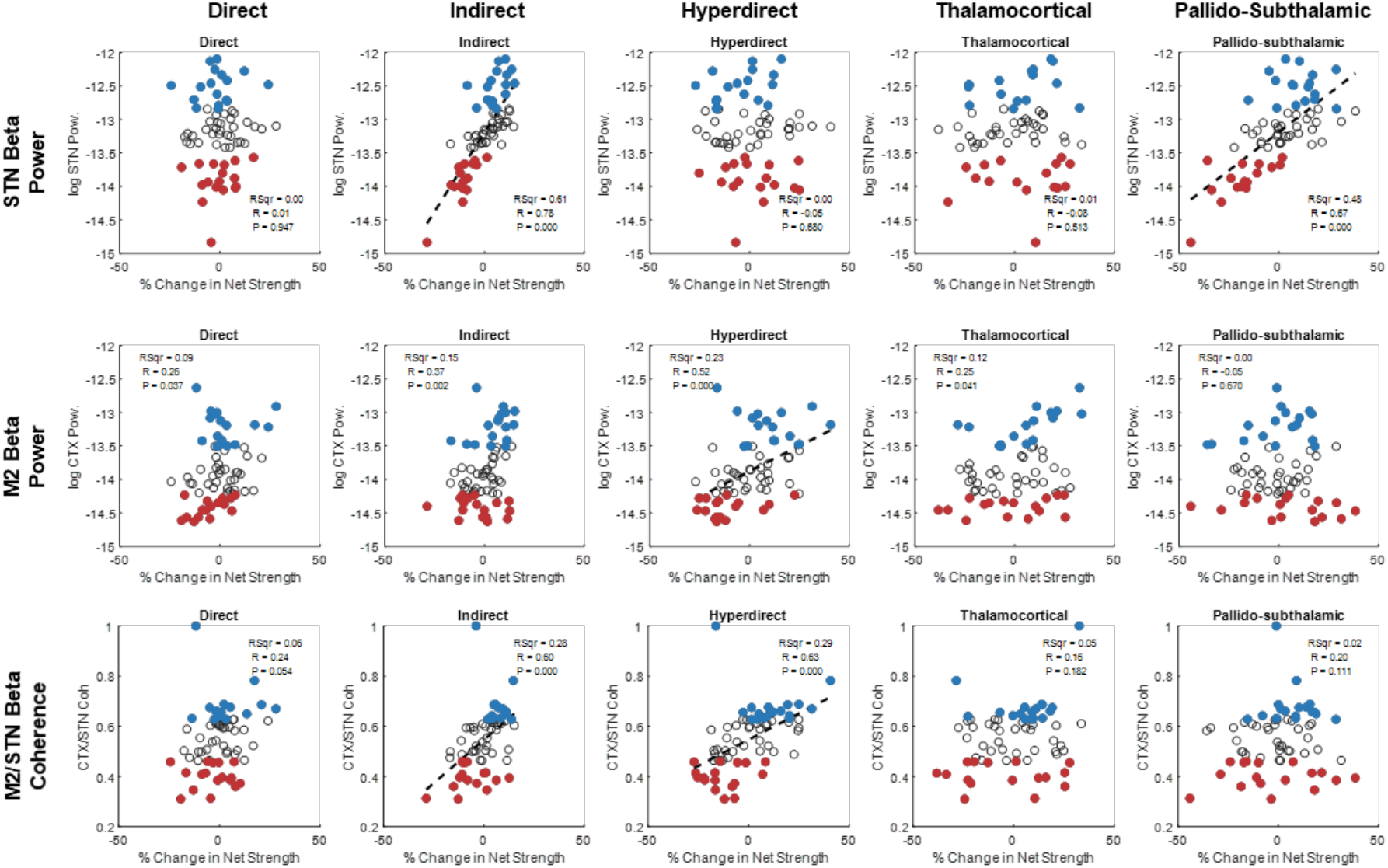
Full set of correlations of beta (14-30 Hz) power in the STN (first row), M2 (second row) and STN/M2 coherence following changes in loop strength. The fitted “Net Strength” in the base model is denoted as 0%. Samples in the upper and lower quartiles of beta power are marked in blue and red respectively. Where there was significant correlation (Spearman’s coefficient; Bonferroni corrected α* = α/n; where n = 15 separate tests) a linear regression is plot and resulting statistics inset.

**Supplementary Information III.**
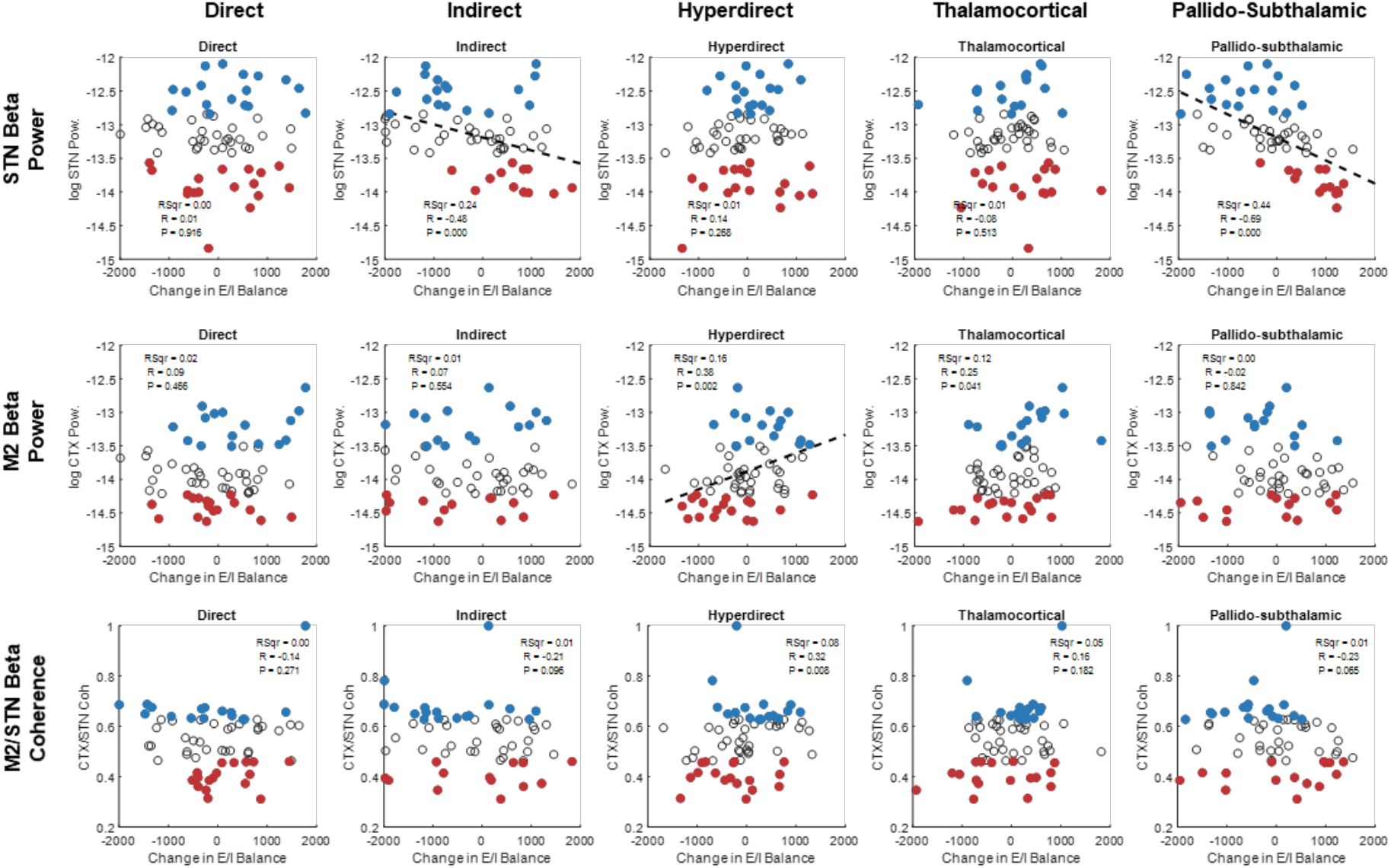
Full set of correlations of beta (14-30 Hz) power in the STN (first row), M2 (second row) and STN/M2 coherence following changes in loop E/I balance. The fitted E/I balance in the base model is denoted as 0%. Samples in the upper and lower quartiles of beta power are marked in blue and red respectively. Where there was significant correlation (Spearman’s coefficient; Bonferroni corrected α* = α/n; where n = 15 separate tests) a linear regression is plot and resulting statistics inset.

**Supplementary Information IV.**
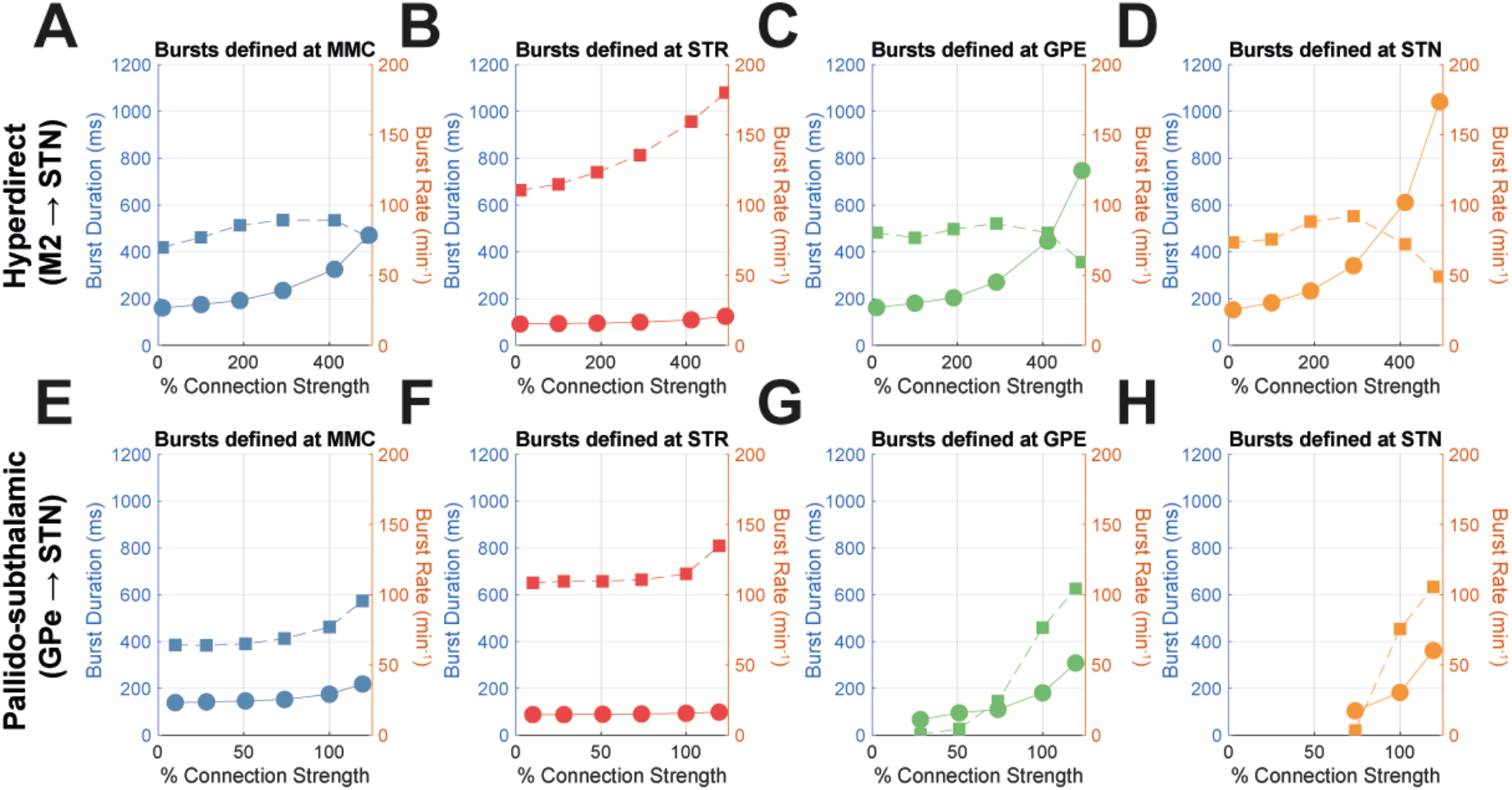
Changes in burst properties associated with changes in connection strength. **(A-D)** Changes relating to modulation of the hyperdirect pathway. **(E-H)** Changes relating the modulation of the pallido-subthalamic pathway.

**Supplementary Information V.**
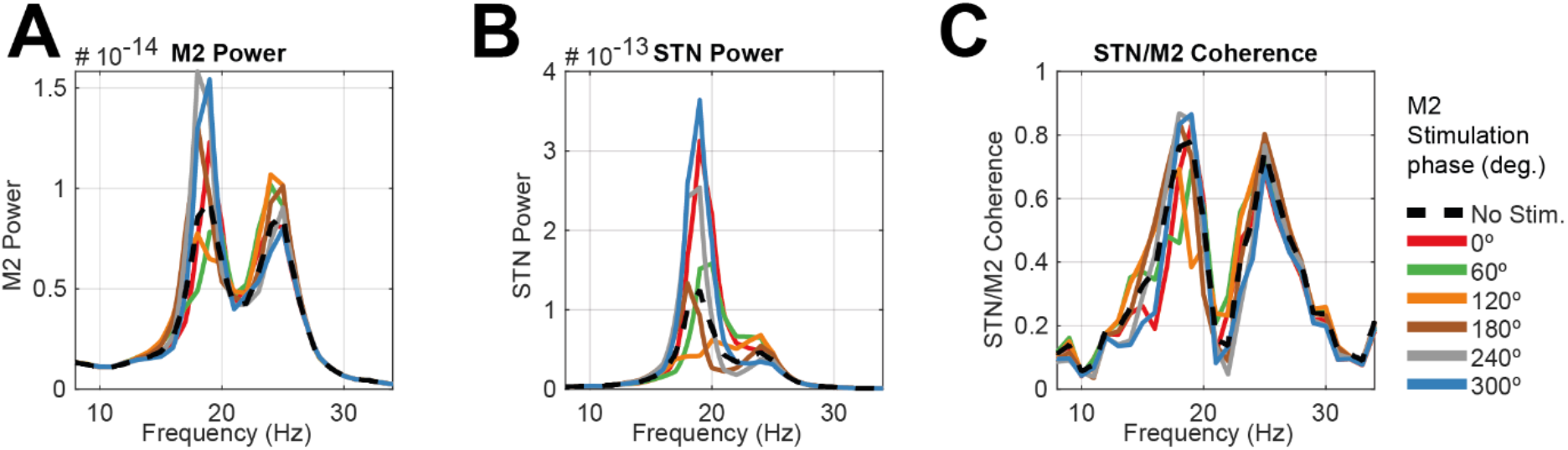
Resulting power spectra from phase locked stimulation of M2 when sensing activity in the STN. **(A)** Power in the cortex, **(B)** Power in the STN, and **(C)** STN/M2 coherence. The angle of stimulation is given by the legend (inset).

**Supplementary Information VI.**
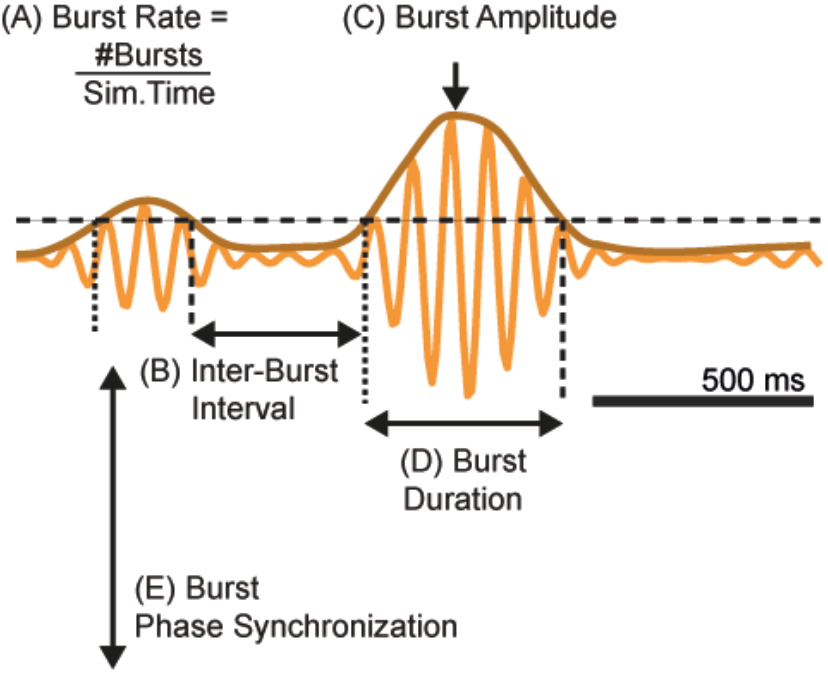
Schematic of the analysis of burst properties. Burst rate (A) is calculated as the total number of bursts within the simulated time, (B) the interburst interval is the time between burst offset and onset, (C) the burst amplitude is the maximum of the within-burst envelope, and (D) the burst duration is the time from onset to offset.

**Supplementary Table I.**

| Table of Model Parameters |  |  |  |  |
| --- | --- | --- | --- | --- |
| **Cortex** | *Prior* | *Prior variance* | *Posterior -log scaling* | *Posterior* |
| Time constant (*ms*) |  |  |  |  |
| *middle (mp)* | 3 | 1/4 | -0.48 | 2 |
| *superficial (sp)* | 2 | 1/4 | -0.27 | 2 |
| *Interneurons (ii)* | 12 | 1/4 | 0.35 | 17 |
| *deep (dp)* | 18 | 1/4 | -0.19 | 15 |
| Synaptic gain (*Hz*) |  |  |  |  |
| *mp* → *mp* (*self inh.*) | -400 | 1/4 | 0.15 | -464.90 |
| *mp* → *sp* | 800 | 1/4 | 0.41 | 1207.82 |
| *ii* → *mp* | -400 | 1/4 | 0.07 | -431.02 |
| *ii* → *ii* (*self inh.*) | -400 | 1/4 | 0.72 | -822.30 |
| *mp* → *ii* | 400 | 1/4 | -0.95 | 155.41 |
| *dp* → *ii* | 400 | 1/4 | 0.35 | 565.31 |
| *sp* → *sp* (*self inh.*) | -400 | 1/4 | 0.01 | -402.60 |
| *sp* → *mp* | 400 | 1/4 | 0.40 | 593.93 |
| *ii* → *dp* | -400 | 1/4 | -0.50 | -241.61 |
| *dp* → *dp* (*self inh.*) | -400 | 1/4 | 0.51 | -667.42 |
| *sp* → *dp* | 800 | 1/4 | -0.92 | 318.44 |
| *ii* → *sp* | -400 | 1/4 | 0.46 | -632.43 |
| *sp* → *ii* | -400 | 1/4 | 0.33 | -554.68 |
| *dp* → *sp* | 400 | 1/4 | -0.07 | 371.34 |
| Input gain (*scalar*) |  |  |  |  |
|  | 1 | 1/16 | -0.066 | 0.9364 |
| **Striatum** | *Prior* | *Prior variance* | *Posterior -log scaling* | *Posterior* |
| Time constant (*ms*) |  |  |  |  |
|  | 8 | 1/8 | -0.05 | 8 |
| Synaptic gain (*Hz*) |  |  |  |  |
| *self inh.* | 400 | 1/8 | 2.12 | 849 |
| Input gain (*scalar*) |  |  |  |  |
|  | 1 | 1/16 | 0.25 | 1.288 |
| **GPe** | *Prior* | *Prior variance* | *Posterior -log scaling* | *Posterior* |
| Time constant (*ms*) |  |  |  |  |
|  | 8 | 1/8 | 0.49 | 13 |
| Input gain (*scalar*) |  |  |  |  |
|  | 1 | 1/16 | -0.207 | 0.813 |
| **STN** | *Prior* | *Prior variance* | *Posterior -log scaling* | *Posterior* |
| Time constant (*ms*) |  |  |  |  |
|  | 4 | 1/8 | -0.68 | 2 |
| Input gain (*scalar*) |  |  |  |  |
|  | 1 | 1/16 | -0.188 | 0.829 |
| **GPI** | *Prior* | *Prior variance* | *Posterior -log scaling* | *Posterior* |
| Time constant (*ms*) |  |  |  |  |
|  | 8 | 1/8 | -0.01 | 8 |
| Input gain (*scalar*) |  |  |  |  |
|  | 1 | 1/16 | -0.14 | 0.874 |
| **Thalamus** | *Prior* | *Prior variance* | *Posterior -log scaling* | *Posterior* |
| Time constant (*ms*) |  |  |  |  |
|  | 8 | 1/8 | 1 | 8 |
| Synaptic gain (*Hz*) |  |  |  |  |
| *self inh.* | -400 | 1/8 | 1.82 | -729 |
| Input gain (*scalar*) |  |  |  |  |
|  | 1 | 1/16 | 0.37 | 1.45 |
| **Extrinsic Parameters** | *Prior* | *Prior variance* | *Posterior -log scaling* | *Posterior* |
| Connection weights (*Hz*) |  |  |  |  |
| M2 → STR | 1600 | 1/4 | -0.22 | 1284 |
| M2 → STN | 2000 | 1/4 | -0.64 | 1055 |
| M2 → Thal | 1000 | 1/4 | 0.164 | 1178 |
| STR → GPe | -2000 | 1/4 | -0.17 | -1687 |
| STR → GPi | -1600 | 1/4 | 0.87 | -3819 |
| GPe → STN | -2000 | 1/4 | 0.9 | -4919 |
| STN → GPe | 2000 | 1/4 | -0.7 | 993 |
| STN → GPi | 1600 | 1/4 | -0.22 | 1284 |
| GPi → Thal | -1000 | 1/4 | 0.54 | -1716 |
| Thal → M2 | 2000 | 1/4 | 0.14 | 2301 |
| Delays (*ms*) |  |  |  |  |
| M2 → STR | 3 | 1/16 | 0.68 | 6 |
| M2 → STN | 3 | 1/16 | -0.03 | 3 |
| M2 → Thal. | 4 | 1/16 | 0.61 | 7.5 |
| STR → GPe | 7 | 1/16 | -0.44 | 5 |
| STR → GPi | 12 | 1/16 | -0.55 | 7 |
| GPe → STN | 1 | 1/16 | -0.58 | 1 |
| STN → GPe | 3 | 1/16 | -0.56 | 2 |
| STN → GPi | 3 | 1/16 | 0.39 | 4 |
| GPi → Thal | 3 | 1/16 | 0.56 | 5.5 |
| Thal → M2 | 8 | 1/16 | 0.65 | 15.5 |

**Supplementary Table II.** We thank all authors and contributors of the toolboxes below:

| **Toolbox Name** | **Author** | **Year** | **Source/Reference** |
| --- | --- | --- | --- |
| **allcomb** | ‘Jos’ | 2018 | https://uk.mathworks.com/matlabcentral/fileexchange/10064-allcomb-varargin |
| **boundedline-pkg** | Kelly Kearney | 2015 | https://github.com/kakearney/boundedline-pkg |
| **brewermap** | Stephen Cobeldick | 2014 | https://github.com/DrosteEffect/BrewerMap |
| **Fieldtrip** | Donders Institute, Radboud University | 2020 | www.fieldtriptoolbox.org/ |
| **linspecer** | Jonathan C. Lansey | 2015 | https://github.com/davidkun/linspecer |
| **neurospec 2.2** | David Halliday | 2018 | http://www.neurospec.org/ |
| **ParforProgMon** | Dylan Muir, Willem-Jan de Goeij, The MathWorks, Inc. | 2016 | https://github.com/DylanMuir/ParforProgMon |
| **splitvec** | Bruno Luong | 2009 | https://uk.mathworks.com/matlabcentral/fileexchange/24255-splitvec |
| **SPM 12** | Wellcome Centre for Human Neuroimaging, University College London | 2020 | https://www.fil.ion.ucl.ac.uk/spm/ |

## References

Ali MM, Sellers KK, Fröhlich F (2013) Transcranial alternating current stimulation modulates large-scale cortical network activity by network resonance. J Neurosci 33:11262–11275.

Amadeus Steiner L, Barreda Tomás FJ, Planert H, Alle H, Vida I, Geiger JRP (2019) Connectivity and dynamics underlying synaptic control of the subthalamic nucleus. J Neurosci 39:2470–2481.

Arlotti M, Rosa M, Marceglia S, Barbieri S, Priori A (2016) The adaptive deep brain stimulation challenge. Parkinsonism Relat Disord 28:12–17.

Baaske MK, Kormann E, Holt AB, Gulberti A, McNamara CG, Pötter-Nerger M, Westphal M, Engel AK, Hamel W, Brown P, Moll CKE, Sharott A (2019) Parkinson’s disease uncovers an underlying sensitivity of subthalamic nucleus neurons to beta-frequency cortical input. bioRxiv:513234.

Baker AP, Brookes MJ, Rezek IA, Smith SM, Behrens T, Probert Smith PJ, Woolrich M (2014) Fast transient networks in spontaneous human brain activity. Elife 3.

Beaumont MA, Zhang W, Balding DJ (2002) Approximate Bayesian Computation in Population Genetics. Genetics 162:2025 LP – 2035.

Bergmann TO, Karabanov A, Hartwigsen G, Thielscher A, Siebner HR (2016) Combining non-invasive transcranial brain stimulation with neuroimaging and electrophysiology: Current approaches and future perspectives. Neuroimage 140:4–19.

Bevan MD, Magill PJ, Terman D, Bolam JP, Wilson CJ (2002) Move to the rhythm: oscillations in the subthalamic nucleus–external globus pallidus network. Trends Neurosci 25:525–531.

Bhatt MB, Bowen S, Rossiter HE, Dupont-Hadwen J, Moran RJ, Friston KJ, Ward NS (2016) Computational modelling of movement-related beta-oscillatory dynamics in human motor cortex. Neuroimage 133:224–232.

Blenkinsop A, Anderson S, Gurney K (2017) Frequency and function in the basal ganglia: the origins of beta and gamma band activity. J Physiol 595:4525–4548.

Bouthour W, Mégevand P, Donoghue J, Lüscher C, Birbaumer N, Krack P (2019) Biomarkers for closed-loop deep brain stimulation in Parkinson disease and beyond. Nat Rev Neurol 15:343–352.

Brazhnik E, McCoy AJ, Novikov N, Hatch CE, Walters JR (2016) Ventral medial thalamic nucleus promotes synchronization of increased high beta oscillatory activity in the basal ganglia-thalamocortical network of the hemiparkinsonian rat. J Neurosci 36:4196–4208.

Brazhnik E, Novikov N, McCoy AJ, Cruz A V, Walters JR (2014) Functional correlates of exaggerated oscillatory activity in basal ganglia output in hemiparkinsonian rats. Exp Neurol 261:563–577.

Bressler SL, Menon V (2010) Large-scale brain networks in cognition: emerging methods and principles. Trends Cogn Sci 14:277–290.

Brittain J-S, Brown P (2014) Oscillations and the basal ganglia: motor control and beyond. Neuroimage 85:637–647.

Brittain JS, Probert-Smith P, Aziz TZ, Brown P (2013) Tremor suppression by rhythmic transcranial current stimulation. Curr Biol 23:436–440.

Brown P (2007) Abnormal oscillatory synchronisation in the motor system leads to impaired movement. Curr Opin Neurobiol 17:656–664.

Brown P, Oliviero A, Mazzone P, Insola A, Tonali P, Di Lazzaro V (2001) Dopamine Dependency of Oscillations between Subthalamic Nucleus and Pallidum in Parkinson’s Disease. J Neurosci 21:1033–1038.

Cagnan H, Denison T, McIntyre C, Brown P (2019a) Emerging technologies for improved deep brain stimulation. Nat Biotechnol 37:1024–1033.

Cagnan H, Duff EP, Brown P (2015) The relative phases of basal ganglia activities dynamically shape effective connectivity in Parkinson’s disease. Brain 138:1667–1678.

Cagnan H, Mallet N, Moll CKE, Gulberti A, Holt AB, Westphal M, Gerloff C, Engel AK, Hamel W, Magill PJ, Brown P, Sharott A (2019b) Temporal evolution of beta bursts in the parkinsonian cortical and basal ganglia network. Proc Natl Acad Sci U S A:201819975.

Cagnan H, Pedrosa D, Little S, Pogosyan A, Cheeran B, Aziz T, Green A, Fitzgerald J, Foltynie T, Limousin P, Zrinzo L, Hariz M, Friston KJ, Denison T, Brown P (2017) Stimulating at the right time: phase-specific deep brain stimulation. Brain 140:132–145.

Castrioto A, Lhommée E, Moro E, Krack P (2014) Mood and behavioural effects of subthalamic stimulation in Parkinson’s disease. Lancet Neurol 13:287–305.

Chu HY, McIver EL, Kovaleski RF, Atherton JF, Bevan MD (2017) Loss of Hyperdirect Pathway Cortico-Subthalamic Inputs Following Degeneration of Midbrain Dopamine Neurons. Neuron 95:1306–1318.e5.

Corbit VL, Whalen TC, Zitelli KT, Crilly SY, Rubin JE, Gittis AH (2016) Pallidostriatal Projections Promote β Oscillations in a Dopamine-Depleted Biophysical Network Model. J Neurosci 36:5556–5571.

Cordon I, Nicolás MJ, Arrieta S, Alegre M, Artieda J, Valencia M (2018) Theta-phase closed-loop stimulation induces motor paradoxical responses in the rat model of Parkinson disease. Brain Stimul 11:231–238.

Crompe B de la, Aristieta A, Leblois A, Elsherbiny S, Boraud T, Mallet N (2020) The globus pallidus orchestrates abnormal network dynamics in a model of Parkinsonism. Nat Commun 11:1–14.

Cruz A V., Mallet N, Magill PJ, Brown P, Averbeck BB (2011) Effects of dopamine depletion on information flow between the subthalamic nucleus and external globus pallidus. 106:2012–2023.

David O, Friston KJ (2003) A neural mass model for MEG/EEG: Neuroimage 20:1743–1755.

de Solages C, Hill BC, Koop MM, Henderson JM, Bronte-Stewart H (2010) Bilateral symmetry and coherence of subthalamic nuclei beta band activity in Parkinson’s disease. Exp Neurol 221:260–266.

Deffains M, Iskhakova L, Katabi S, Israel Z, Bergman H (2018) Longer β oscillatory episodes reliably identify pathological subthalamic activity in Parkinsonism. Mov Disord 33:1609–1618.

Delaville C, Cruz A V, McCoy AJ, Brazhnik E, Avila I, Novikov N, Walters JR (2014) Oscillatory Activity in Basal Ganglia and Motor Cortex in an Awake Behaving Rodent Model of Parkinson’s Disease. Basal Ganglia 3:221–227.

Dostrovsky J, Bergman H (2004) Oscillatory activity in the basal ganglia--relationship to normal physiology and pathophysiology. Brain 127:721–722.

Engel AK, Fries P (2010) Beta-band oscillations—signalling the status quo? Curr Opin Neurobiol 20:156–165.

Farries MA, Kita H, Wilson CJ (2010) Dynamic spike threshold and zero membrane slope conductance shape the response of subthalamic neurons to cortical input. J Neurosci 30:13180–13191.

Feingold J, Gibson DJ, DePasquale B, Graybiel AM (2015) Bursts of beta oscillation differentiate postperformance activity in the striatum and motor cortex of monkeys performing movement tasks. Proc Natl Acad Sci 112:13687–13692.

Fogelson N, Williams D, Tijssen M, van Bruggen G, Speelman H, Brown P (2006) Different Functional Loops between Cerebral Cortex and the Subthalmic Area in Parkinson’s Disease. Cereb Cortex 16:64–75.

Fountas Z, Shanahan M (2017) The role of cortical oscillations in a spiking neural network model of the basal ganglia. PLoS One 12:e0189109.

Frank MJ, Samanta J, Moustafa AA, Sherman SJ (2007) Hold your horses: impulsivity, deep brain stimulation, and medication in parkinsonism. Science 318:1309–1312.

Gillies A, Willshaw D (2004) Models of the subthalamic nucleus: The importance of intranuclear connectivity. Med Eng Phys 26:723–732.

Grado LL, Johnson MD, Netoff TI (2018) Bayesian adaptive dual control of deep brain stimulation in a computational model of Parkinson’s disease Santaniello S, ed. PLOS Comput Biol 14:e1006606.

Hahn G, Bujan AF, Frégnac Y, Aertsen A, Kumar A (2014) Communication through Resonance in Spiking Neuronal Networks Brunel N, ed. PLoS Comput Biol 10:e1003811.

Halliday DM, Senik MH, Stevenson CW, Mason R (2016) Non-parametric directionality analysis – Extension for removal of a single common predictor and application to time series. J Neurosci Methods 268:87–97.

Hammond C, Bergman H, Brown P (2007) Pathological synchronization in Parkinson’s disease: networks, models and treatments. Trends Neurosci 30:357–364.

Hannah R, Muralidharan V, Sundby KK, Aron AR (2019) Temporally-precise disruption of prefrontal cortex informed by the timing of beta bursts impairs human action-stopping. bioRxiv:843557.

Herz DM, Little S, Pedrosa DJ, Tinkhauser G, Cheeran B, Foltynie T, Bogacz R, Brown P (2018) Mechanisms Underlying Decision-Making as Revealed by Deep-Brain Stimulation in Patients with Parkinson’s Disease. Curr Biol 28:1169–1178.e6.

Holt AB, Kormann E, Gulberti A, Pötter-Nerger M, McNamara CG, Cagnan H, Baaske MK, Little S, Köppen JA, Buhmann C, Westphal M, Gerloff C, Engel AK, Brown P, Hamel W, Moll CKE, Sharott A (2019) Phase-Dependent Suppression of Beta Oscillations in Parkinson’s Disease Patients. J Neurosci 39:1119–1134.

Holt AB, Netoff TI (2014) Origins and suppression of oscillations in a computational model of Parkinson’s disease. J Comput Neurosci 37:505–521.

Jahfari S, Waldorp L, van den Wildenberg WPM, Scholte HS, Ridderinkhof KR, Forstmann BU (2011) Effective Connectivity Reveals Important Roles for Both the Hyperdirect (Fronto-Subthalamic) and the Indirect (Fronto-Striatal-Pallidal) Fronto-Basal Ganglia Pathways during Response Inhibition. J Neurosci 31.

Jansen BH, Rit VG (1995) Electroencephalogram and visual evoked potential generation in a mathematical model of coupled cortical columns. Biol Cybern 73:357–366.

Jenkinson N, Brown P (2011) New insights into the relationship between dopamine, beta oscillations and motor function. Trends Neurosci 34:611–618.

Kahan J, Mancini L, Flandin G, White M, Papadaki A, Thornton J, Yousry T, Zrinzo L, Hariz M, Limousin P, Friston K, Foltynie T (2019) Deep brain stimulation has state-dependent effects on motor connectivity in Parkinson’s disease. Brain 142:2417–2431.

Karabanov A, Thielscher A, Siebner HR (2016) Transcranial brain stimulation: Closing the loop between brain and stimulation. Curr Opin Neurol 29:397–404.

Khanna P, Carmena JM (2015) Neural oscillations: beta band activity across motor networks. Curr Opin Neurobiol 32:60–67.

Khanna P, Carmena JM (2017) Beta band oscillations in motor cortex reflect neural population signals that delay movement onset. Elife 6.

Kumar A, Cardanobile S, Rotter S, Aertsen A (2011) The role of inhibition in generating and controlling Parkinson’s disease oscillations in the Basal Ganglia. Front Syst Neurosci 5:86.

Lalo E, Thobois S, Sharott A, Polo G, Mertens P, Pogosyan A, Brown P (2008) Patterns of Bidirectional Communication between Cortex and Basal Ganglia during Movement in Patients with Parkinson Disease. J Neurosci 28.

Leblois A, Boraud T, Meissner W, Bergman H, Hansel D (2006) Competition between Feedback Loops Underlies Normal and Pathological Dynamics in the Basal Ganglia. J Neurosci 26:3567–3583.

Levy R (2002) Dependence of subthalamic nucleus oscillations on movement and dopamine in Parkinson’s disease. Brain 125:1196–1209.

Liepe J, Kirk P, Filippi S, Toni T, Barnes CP, Stumpf MPH (2014) A framework for parameter estimation and model selection from experimental data in systems biology using approximate Bayesian computation. Nat Protoc 9:439–456.

Little S, Beudel M, Zrinzo L, Foltynie T, Limousin P, Hariz M, Neal S, Cheeran B, Cagnan H, Gratwicke J, Aziz TZ, Pogosyan A, Brown P (2016a) Bilateral adaptive deep brain stimulation is effective in Parkinson’s disease. J Neurol Neurosurg Psychiatry 87:717–721.

Little S, Bonaiuto J, Barnes G, Bestmann S (2019) Human motor cortical beta bursts relate to movement planning and response errors Ganguly K, ed. PLOS Biol 17:e3000479.

Little S, Pogosyan A, Neal S, Zavala B, Zrinzo L, Hariz M, Foltynie T, Limousin P, Ashkan K, FitzGerald J, Green AL, Aziz TZ, Brown P (2013) Adaptive deep brain stimulation in advanced Parkinson disease. Ann Neurol 74:449–457.

Little S, Tripoliti E, Beudel M, Pogosyan A, Cagnan H, Herz D, Bestmann S, Aziz T, Cheeran B, Zrinzo L, Hariz M, Hyam J, Limousin P, Foltynie T, Brown P (2016b) Adaptive deep brain stimulation for Parkinson’s disease demonstrates reduced speech side effects compared to conventional stimulation in the acute setting. J Neurol Neurosurg Psychiatry 87:1388–1389.

Litvak V, Jha A, Eusebio A, Oostenveld R, Foltynie T, Limousin P, Zrinzo L, Hariz MI, Friston K, Brown P (2011) Resting oscillatory cortico-subthalamic connectivity in patients with Parkinson’s disease. Brain 134:359–374.

Liu C, Zhu Y, Liu F, Wang J, Li H, Deng B, Fietkiewicz C, Loparo KA (2017) Neural mass models describing possible origin of the excessive beta oscillations correlated with Parkinsonian state. Neural Networks 88:65–73.

Llinás RR, Ribary U, Jeanmonod D, Kronberg E, Mitra PP (1999) Thalamocortical dysrhythmia: A neurological and neuropsychiatric syndrome characterized by magnetoencephalography. Proc Natl Acad Sci U S A 96:15222–15227.

Lundqvist M, Rose J, Herman P, Brincat SL, Buschman TJ, Miller EK (2016) Gamma and Beta Bursts Underlie Working Memory. Neuron 90:152–164.

Magill PJ, Pogosyan A, Sharott A, Csicsvari J, Bolam JP, Brown P (2006) Changes in functional connectivity within the rat striatopallidal axis during global brain activation in vivo. J Neurosci 26:6318–6329.

Magill PJ, Sharott A, Bolam JP, Brown P (2004) Brain State–Dependency of Coherent Oscillatory Activity in the Cerebral Cortex and Basal Ganglia of the Rat. J Neurophysiol 92:2122–2136.

Mallet N, Pogosyan A, Marton LF, Bolam JP, Brown P, Magill PJ (2008a) Parkinsonian Beta Oscillations in the External Globus Pallidus and Their Relationship with Subthalamic Nucleus Activity. J Neurosci 28:14245–14258.

Mallet N, Pogosyan A, Sharott A, Csicsvari J, Bolam JP, Brown P, Magill PJ (2008b) Disrupted Dopamine Transmission and the Emergence of Exaggerated Beta Oscillations in Subthalamic Nucleus and Cerebral Cortex. J Neurosci 28:4795–4806.

Marreiros AC, Cagnan H, Moran RJ, Friston KJ, Brown P (2013) Basal ganglia–cortical interactions in Parkinsonian patients. Neuroimage 66:301–310.

Martin DM, Liu R, Alonzo A, Green M, Loo CK (2014) Use of transcranial direct current stimulation (tDCS) to enhance cognitive training: effect of timing of stimulation. Exp Brain Res 232:3345–3351.

Mathai A, Ma Y, Paré J-F, Villalba RM, Wichmann T, Smith Y (2015) Reduced cortical innervation of the subthalamic nucleus in MPTP-treated parkinsonian monkeys. Brain 138:946–962.

McCarthy MM, Moore-Kochlacs C, Gu X, Boyden ES, Han X, Kopell N (2011) Striatal origin of the pathologic beta oscillations in Parkinson’s disease. Proc Natl Acad Sci U S A 108:11620–11625.

McIntyre CC, Hahn PJ (2010) Network perspectives on the mechanisms of deep brain stimulation. Neurobiol Dis 38:329–337.

Mirzaei A, Kumar A, Leventhal D, Mallet N, Aertsen A, Berke J, Schmidt R (2017) Sensorimotor Processing in the Basal Ganglia Leads to Transient Beta Oscillations during Behavior. J Neurosci 37:11220–11232.

Moran RJ, Mallet N, Litvak V, Dolan RJ, Magill PJ, Friston KJ, Brown P (2011) Alterations in brain connectivity underlying beta oscillations in parkinsonism Kording KP, ed. PLoS Comput Biol 7:e1002124.

Palmer C et al. (2016) A New Framework to Explain Sensorimotor Beta Oscillations. Trends Cogn Sci 20:321–323.

Pavlides A, Hogan SJ, Bogacz R (2015) Computational Models Describing Possible Mechanisms for Generation of Excessive Beta Oscillations in Parkinson’s Disease. PLOS Comput Biol 11:e1004609.

Paxinos G, Watson C (2007) The rat brain in stereotaxic coordinates. Elsevier.

Peles O, Werner-Reiss U, Bergman H, Israel Z, Vaadia E (2020) Phase-Specific Microstimulation Differentially Modulates Beta Oscillations and Affects Behavior. Cell Rep 30:2555–2566.e3.

Pirulli C, Fertonani A, Miniussi C (2013) The role of timing in the induction of neuromodulation in perceptual learning by transcranial electric stimulation. Brain Stimul 6:683–689.

Polanía R, Nitsche MA, Korman C, Batsikadze G, Paulus W (2012) The importance of timing in segregated theta phase-coupling for cognitive performance. Curr Biol 22:1314–1318.

Reis C, Sharott A, Magill PJ, van Wijk BCM, Parr T, Zeidman P, Friston KJ, Cagnan H (2019) Thalamocortical dynamics underlying spontaneous transitions in beta power in Parkinsonism. Neuroimage 193:103–114.

Rosa M, Arlotti M, Ardolino G, Cogiamanian F, Marceglia S, Di Fonzo A, Cortese F, Rampini PM, Priori A (2015) Adaptive deep brain stimulation in a freely moving parkinsonian patient. Mov Disord 30:1003–1005.

Rosenblum M, Pikovsky A (2004) Delayed feedback control of collective synchrony: An approach to suppression of pathological brain rhythms. Phys Rev E – Stat Physics, Plasmas, Fluids, Relat Interdiscip Top 70:11.

Sanabria DE, Johnson LA, Yu Y, Busby Z, Nebeck S, Zhang J, Harel N, Johnson MD, Molnar GF, Vitek JL (2020) Suppressing and amplifying neural oscillations in real-time using phase-locked electrical stimulation: concept, optimization and in-vivo testing. bioRxiv:2020.02.09.940643.

Sanders TH, Jaeger D (2016) Optogenetic stimulation of cortico-subthalamic projections is sufficient to ameliorate bradykinesia in 6-ohda lesioned mice. Neurobiol Dis 95:225–237.

Schnitzler A, Gross J (2005) Normal and pathological oscillatory communication in the brain. Nat Rev Neurosci 6:285–296.

Sharott A, Gulberti A, Hamel W, Köppen JA, Münchau A, Buhmann C, Pötter-Nerger M, Westphal M, Gerloff C, Moll CKE, Engel AK (2018) Spatio-temporal dynamics of cortical drive to human subthalamic nucleus neurons in Parkinson’s disease. Neurobiol Dis 112:49–62.

Sharott A, Magill PJ, Bolam JP, Brown P (2005a) Directional analysis of coherent oscillatory field potentials in the cerebral cortex and basal ganglia of the rat. J Physiol 562:951–963.

Sharott A, Magill PJ, Harnack D, Kupsch A, Meissner W, Brown P (2005b) Dopamine depletion increases the power and coherence of beta-oscillations in the cerebral cortex and subthalamic nucleus of the awake rat. Eur J Neurosci 21:1413–1422.

Sharott A, Vinciati F, Nakamura KC, Magill PJ (2017) A Population of Indirect Pathway Striatal Projection Neurons Is Selectively Entrained to Parkinsonian Beta Oscillations. J Neurosci 37:9977–9998.

Sherman MA, Lee S, Law R, Haegens S, Thorn CA, Hämäläinen MS, Moore CI, Jones SR (2016) Neural mechanisms of transient neocortical beta rhythms: Converging evidence from humans, computational modeling, monkeys, and mice. Proc Natl Acad Sci U S A 113:E4885–94.

Shin H, Law R, Tsutsui S, Moore CI, Jones SR (2017) The rate of transient beta frequency events predicts behavior across tasks and species. Elife 6.

Shouno O, Tachibana Y, Nambu A, Doya K (2017) Computational Model of Recurrent Subthalamo-Pallidal Circuit for Generation of Parkinsonian Oscillations. Front Neuroanat 11:21.

Siegle JH, Wilson MA (2014) Enhancement of encoding and retrieval functions through theta phase-specific manipulation of hippocampus. Elife 3.

Silvanto J, Muggleton N, Walsh V (2008) State-dependency in brain stimulation studies of perception and cognition. Trends Cogn Sci 12:447–454.

Stagg CJ, Jayaram G, Pastor D, Kincses ZT, Matthews PM, Johansen-Berg H (2011) Polarity and timing-dependent effects of transcranial direct current stimulation in explicit motor learning. Neuropsychologia 49:800–804.

Steriade M (2000) Corticothalamic resonance, states of vigilance and mentation. Neuroscience 101:243–276.

Swann NC, de Hemptinne C, Thompson MC, Miocinovic S, Miller AM, Gilron R, Ostrem JL, Chizeck HJ, Starr PA (2018) Adaptive deep brain stimulation for Parkinson’s disease using motor cortex sensing. J Neural Eng 15:46006.

Swann NC, Hemptinne C de, Miocinovic S, Qasim S, Ostrem JL, Galifianakis NB, Luciano MS, Wang SS, Ziman N, Taylor R, Starr PA (2017) Chronic multisite brain recordings from a totally implantable bidirectional neural interface: experience in 5 patients with Parkinson’s disease. J Neurosurg JNS 128:605–616.

Tachibana Y, Iwamuro H, Kita H, Takada M, Nambu A (2011) Subthalamo-pallidal interactions underlying parkinsonian neuronal oscillations in the primate basal ganglia. Eur J Neurosci 34:1470–1484.

Tass PA (2000) Stochastic Phase Resetting: A Theory for Deep Brain Stimulation. Prog Theor Phys Suppl 139:301–313.

Terman D, Rubin JE, Yew AC, Wilson CJ (2002) Activity patterns in a model for the subthalamopallidal network of the basal ganglia. J Neurosci 22:2963–2976.

Thut G, Miniussi C, Gross J (2012) The functional importance of rhythmic activity in the brain. Curr Biol 22:R658–63.

Tinkhauser G, Pogosyan A, Little S, Beudel M, Herz DM, Tan H, Brown P (2017a) The modulatory effect of adaptive deep brain stimulation on beta bursts in Parkinson’s disease. Brain 140:1053–1067.

Tinkhauser G, Pogosyan A, Tan H, Herz DM, Kühn AA, Brown P (2017b) Beta burst dynamics in Parkinson’s disease OFF and ON dopaminergic medication. Brain 140:2968–2981.

Tinkhauser G, Torrecillos F, Duclos Y, Tan H, Pogosyan A, Fischer P, Carron R, Welter M-L, Karachi C, Vandenberghe W, Nuttin B, Witjas T, Régis J, Azulay J-P, Eusebio A, Brown P (2018) Beta burst coupling across the motor circuit in Parkinson’s disease. Neurobiol Dis 117:217–225.

Torrecillos F, Tinkhauser G, Fischer P, Green AL, Aziz TZ, Foltynie T, Limousin P, Zrinzo L, Ashkan K, Brown P, Tan H (2018) Modulation of Beta Bursts in the Subthalamic Nucleus Predicts Motor Performance. J Neurosci 38:8905–8917.

Uhlhaas PJ, Singer W (2006) Neural Synchrony in Brain Disorders: Relevance for Cognitive Dysfunctions and Pathophysiology. Neuron 52:155–168.

van Albada SJ, Gray RT, Drysdale PM, Robinson PA (2009) Mean-field modeling of the basal ganglia-thalamocortical system. II. Dynamics of parkinsonian oscillations. J Theor Biol 257:664–688.

van Wijk BCM, Beek PJ, Daffertshofer A (2012) Neural synchrony within the motor system: what have we learned so far? Front Hum Neurosci 6:252.

van Wijk BCM, Cagnan H, Litvak V, Kühn AA, Friston KJ (2018) Generic dynamic causal modelling: An illustrative application to Parkinson’s disease. Neuroimage 181:818–830.

Volkmann J et al. (2009) Long-term effects of pallidal or subthalamic deep brain stimulation on quality of life in Parkinson’s disease. Mov Disord 24:1154–1161.

Wang DD, de Hemptinne C, Miocinovic S, Ostrem JL, Galifianakis NB, San Luciano M, Starr PA (2018) Pallidal Deep-Brain Stimulation Disrupts Pallidal Beta Oscillations and Coherence with Primary Motor Cortex in Parkinson’s Disease. J Neurosci 38:4556–4568.

Weerasinghe G, Duchet B, Cagnan H, Brown P, Bick C, Bogacz R (2019) Predicting the effects of deep brain stimulation using a reduced coupled oscillator model. PLoS Comput Biol 15:e1006575.

Weinberger M, Mahant N, Hutchison WD, Lozano AM, Moro E, Hodaie M, Lang AE, Dostrovsky JO (2006) Beta oscillatory activity in the subthalamic nucleus and its relation to dopaminergic response in Parkinson’s disease. J Neurophysiol 96:3248–3256.

West T, Farmer S, Berthouze L, Jha A, Beudel M, Foltynie T, Limousin P, Zrinzo L, Brown P, Litvak V (2016) The Parkinsonian Subthalamic Network: Measures of Power, Linear, and Non-linear Synchronization and their Relationship to L-DOPA Treatment and OFF State Motor Severity. Front Hum Neurosci 10:517.

West TO, Berthouze L, Farmer SF, Cagnan H, Litvak V (2019) Mechanistic Inference of Brain Network Dynamics with Approximate Bayesian Computation. bioRxiv:785568.

West TO, Berthouze L, Halliday DM, Litvak V, Sharott A, Magill PJ, Farmer SF (2018) Propagation of Beta/Gamma Rhythms in the Cortico-Basal Ganglia Circuits of the Parkinsonian Rat. J Neurophysiol:jn.00629.2017.

West TO, Halliday DM, Bressler SL, Farmer SF, Litvak V (2020) Measuring Directed Functional Connectivity Using Non-Parametric Directionality Analysis: Validation and Comparison with Non-Parametric Granger Causality. Neuroimage:116796.

Williams D, Tijssen M, Van Bruggen G, Bosch A, Insola A, Di Lazzaro V, Mazzone P, Oliviero A, Quartarone A, Speelman H, Brown P (2002) Dopamine-dependent changes in the functional connectivity between basal ganglia and cerebral cortex in humans. Brain 125:1558–1569.

Wilson D, Moehlis J (2015) Clustered Desynchronization from High-Frequency Deep Brain Stimulation. PLoS Comput Biol 11:1–26.

Witt A, Palmigiano A, Neef A, El Hady A, Wolf F, Battaglia D (2013) Controlling the oscillation phase through precisely timed closed-loop optogenetic stimulation: a computational study. Front Neural Circuits 7:49.

Zanos S, Rembado I, Chen D, Fetz EE (2018) Phase-Locked Stimulation during Cortical Beta Oscillations Produces Bidirectional Synaptic Plasticity in Awake Monkeys. Curr Biol 28:2515–2526.e4.

